# Multi-component functionalized *Bifidobacterium longum* hydrogel for multi-target integrated therapy of colitis-associated anxiety and depression

**DOI:** 10.64898/2026.03.10.710940

**Authors:** Shuo Zhang, Yujie Zhang, Jiansheng He, Shunlian Li, Qingyan Ma, Qiao Li, Yudan Zhang, Yiyang Wang, Shaobo Ma, Songyan Jin, Chune Li, Xueyong Xie, Hang Zhang, Junze Deng, Xueqin Song, Daocheng Wu, Xiancang Ma, Feng Zhu

**Author notes:** These authors contributed equally to this work (Shuo Zhang, Yujie Zhang, Jiansheng He).

## Abstract

Inflammatory bowel diseases (IBDs) are frequently accompanied by anxiety and depression, largely driven by perturbed gut-brain axis signaling. However, current oral therapies remain constrained by the spatial and functional separation between intestinal inflammation and central nervous system dysfunction. Here, we present a comprehensive gut-brain dual region integrated therapeutic strategy based on functionalized *Bifidobacterium longum* hydrogel (INPs@BL@Gel), in which baicalin and tyrosine are coordinated with Fe(III) to form infinite coordination polymers (ICPs), coated with inulin, assembled onto *Bifidobacterium longum* (BL), and subsequently encapsulated within a pH- and matrix metalloproteinase-responsive silk fibroin-gelatin hydrogel. INPs@BL@Gel exhibits high drug-loading, effective gastric protection, inflammation-triggered release, and long-term intestinal colonization. Within the inflamed intestine, BL and components synergistically suppress inflammatory responses, restore gut microbiota homeostasis, and promote intestinal barrier repair through multi-target integrated therapy. Importantly, BL combined with components markedly enhances the production of beneficial neuroactive metabolites such as homovanillic acid and short-chain fatty acids, which integrated regulate neuroinflammation, preserve synaptic function, and facilitate blood-brain barrier repair via the gut-brain axis. In vivo studies demonstrate that INPs@BL@Gel not only exert potent therapeutic efficacy against colitis and effectively alleviate associated depression, but also reshape the gut microbiota and restore barrier integrity, achieving an remarkable comprehensive therapeutic effect.

Scheme 1.
(a) Schematic diagram of the design and preparation of functionalized *Bifidobacterium longum* hydrogel. (b) Exploration of the mechanism of INPs@BL@Gel in treating colitis-associated anxiety and depression through a dual-site multi-target synergistic strategy.

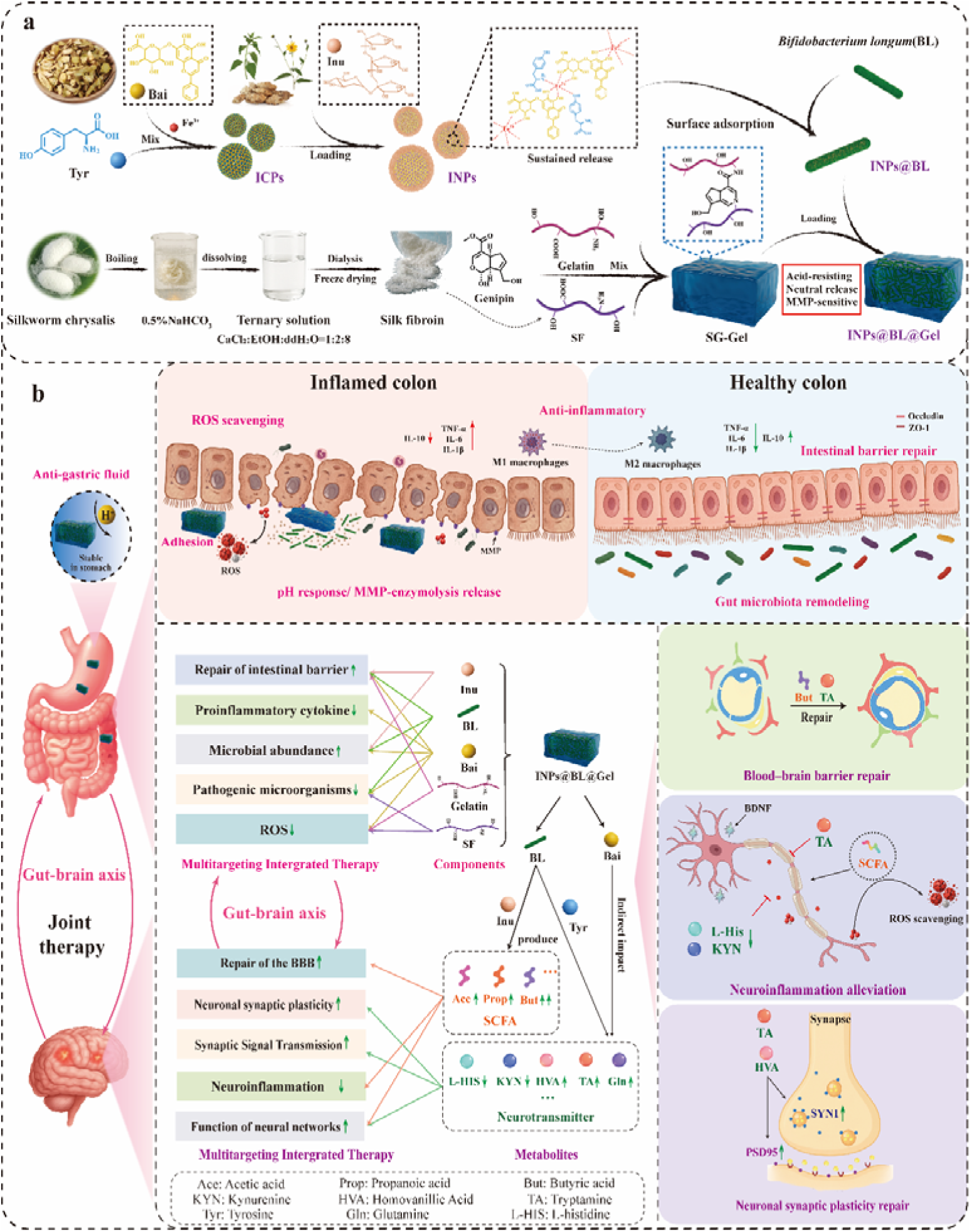

## 1. Introduction

Inflammatory bowel diseases (IBDs) are complex, immune-mediated conditions characterized by chronic, relapsing inflammation of the gastrointestinal tract, and the two major subtypes are Crohn’s disease (CD) and ulcerative colitis (UC).^[1,2]^ In addition to persistent intestinal damage, a growing body of epidemiological and clinical studies demonstrates that individuals with IBDs often experience a higher prevalence of mental health comorbidities, particularly anxiety and depression, compared to the general population and patients with other chronic diseases.^[3–5]^ These mental disorders significantly diminish quality of life and can exacerbate intestinal inflammation through stress-induced pathways, immune dysregulation, and disrupted neuroendocrine responses.^[6–8]^ Collectively, these interactions form a bidirectional loop linking the gut and brain (gut-brain axis), complicating long-term management of IBDs. Thus, there is a clear need for therapeutic strategies that can simultaneously target both IBDs and anxiety and depression associated with colitis.

Currently, oral administration remains the primary treatment method for IBDs and related anxiety and depression due to its safety profile, patient compliance, and convenience.^[9]^ However, existing oral treatment strategies are hindered by the issue of “site separation”. On one hand, anti-inflammatory drugs or microbial preparations primarily address local intestinal inflammation, with limited effects on mental health.^[10,11]^ On the other hand, although antidepressants or anxiolytics can modulate central nervous system (CNS) function, they are insufficient in managing the occurrence and recurrence of intestinal inflammation.^[8,12]^ More importantly, intestinal inflammation and mental disorders are not isolated from each other; instead, they form a highly interconnected, bidirectional regulatory network through the gut-brain axis. This network involves neural pathways (e.g., the vagus nerve and enteric nervous system), immune signals (e.g., the trans-barrier transmission of pro-inflammatory cytokines), endocrine axes (e.g., the hypothalamic-pituitary-adrenal axis), and microbial metabolites (e.g., short-chain fatty acids (SCFAs), neurotransmitter precursors).^[13–15]^ Chronic intestinal inflammation can trigger or worsen CNS dysfunction via these pathways, while mental health conditions can, in turn, intensify the inflammatory response, creating a vicious cycle.^[7,16]^ Furthermore, the dysbiosis of intestinal microbiota and the impairment of the intestinal and blood-brain barrier (BBB) further diminish the long-term efficacy of single-target therapies. Therefore, a therapeutic approach targeting only one organ or pathway is unlikely to effectively control colitis-associated anxiety and depression.

In recent years, probiotics have attracted significant attention for the oral treatment of IBDs due to their favorable biosafety profile and potential to regulate gut microbiota, enhance the mucosal barrier, and restore immune homeostasis.^[17–19]^ To overcome challenges such as degradation in the gastrointestinal tract and limited colonization efficiency, strategies including hydrogel encapsulation, genetic modification, and composite formulations have been developed. Notably, probiotic-based treatments are not limited to local effects on the gut; their metabolites can also influence CNS function through the gut-brain axis, providing a new avenue for the integrated management of colitis-associated depression. For instance, Wang et al. (2022) developed a light-responsive *Lactococcus lactis* system that interacts with the enteric nervous system (ENS). In this system, glucagon-like peptide-1 (GLP-1) signaling through GLP-1 receptors in the ENS activates neuronal nitric oxide synthase (nNOS), stimulating vagal pathways and modulating CNS activity.^[20]^ More recently, Jia et al. (2024) conducted a large cohort study showing that *Bifidobacterium longum* (BL) can synthesize homovanillic acid (HVA) from Tyr. In a mouse model of depression, HVA selectively enhanced the expression of the presynaptic protein synapsin 1 (SYN1) in the hippocampus, providing experimental support for the identification of mechanisms and potential targets involved in colitis-associated depression.^[21]^ These studies offer preliminary validation for the concept of “gut-targeted, brain-modulating” therapies.

Despite these advances, probiotic-based systems still have notable limitations that restrict their ability to coordinate the treatment of colitis-associated depression. (1) Single-dimensional intervention: Most studies focus on alleviating intestinal inflammation, with only indirect effects on mental disorders through microbial metabolites. Rarely is a multi-component, multi-metabolite approach considered, and integrated designs targeting both gut and brain functions are often lacking. As a result, these strategies struggle to break the vicious cycle of “intestinal inflammation-mental disorders”.^[22,23]^ (2) Limited control over metabolites: Many systems rely on the natural metabolism of probiotics, making it difficult to regulate the types and quantities of metabolites produced.^[24,25]^ Additionally, most research focuses on single metabolites, such as HVA or SCFAs, rather than integrating multiple metabolites.^[21,26]^ This limitation restricts the depth and consistency of therapeutic effects on mental disorders. (3) Insufficient engineering integration: Effective treatment of both IBDs and mental disorders often requires multiple components and functions. Integrating these into a single probiotic-based system is highly challenging, as each component must perform multiple functions. The selection and assembly requirements for probiotics and components are extremely high. However, most current probiotic systems are based on simple assembly and have not achieved a unified multi-function system that maintains stability, intestinal colonization, and simultaneous integration of “inflammation suppression, barrier repair, microbiota remodeling, and mental regulation”.^[27–29]^ These limitations hinder the ability of current systems to address the traditional “site separation” problem and provide comprehensive intervention for colitis-associated anxiety and depression.

To address these challenges, this study proposes a comprehensive therapeutic strategy based on “gut-brain dual-region synergistic therapy”. This approach integrates barrier function repair and intestinal microbiota remodeling through a carefully designed probiotic composite system. By employing multi-component, multi-target therapy in gut-brain two sites, this strategy addresses both intestinal inflammation and colitis-associated anxiety and depression, overcoming the limitations of “site separation” in traditional oral treatments. Additionally, the long-term colonization of probiotics supports continuous remodeling of the gut microbiota and sustained repair of the intestinal barrier.^[10,30]^ Meanwhile, a range of key metabolites (e.g., SCFAs and tryptamine molecules) supports BBB function and the homeostasis of the CNS.^[31,32]^ More importantly, a probiotic system with each component serving multiple purposes has been established to achieve the above-mentioned functions.

The strategic design of this study focuses on modulating the types and levels of metabolites by regulating interactions between probiotics and functional components. This enables the targeted enhancement of beneficial metabolites while suppressing potentially harmful ones. BL was selected as the core probiotic strain, given its strong intestinal adaptability and favorable biosafety profile.^[21]^ Studies have shown that BL can repair the intestinal barrier by inhibiting pathogen colonization, enhancing epithelial tight junctions, and regulating immune homeostasis.^[33]^ Moreover, BL utilizes exogenous Tyr to synthesize neuroactive metabolites, including HVA, and can influence the functions of other gut probiotics, indirectly promoting butyric acid and tryptophan production.^[21,33,34]^ Among these metabolites, HVA affects the CNS via the gut-brain axis, playing a role in mood regulation by maintaining synaptic homeostasis and neuroautophagy.^[21]^ Tryptamine has also been linked to reduced neuroinflammation and improved synaptic function.^[35]^ Additionally, this study incorporates the natural products baicalin (Bai) and inulin (Inu) as complementary functional components. Bai has anti-inflammatory and neuroprotective properties, potentially contributing to emotional regulation through modulation of monoamine neurotransmitter pathways.^[36]^ Inu, a prebiotic, promotes selective growth of *Bifidobacterium* and regulates microbial metabolic networks through “cross-feeding”, which further enhances SCFAs production.^[37]^ Butyrate, a key SCFAs, plays a critical role in maintaining both the intestinal barrier and BBB functions.^[38]^ The combination of BL, Bai, Tyr, and Inu is expected to enhance the production of beneficial neuroactive metabolites and SCFAs while suppressing adverse metabolic pathways, leading to multi-target regulation of anxiety and depression.Meanwhile BL and components synergistically suppress inflammatory responses, restore gut microbiota homeostasis, and promote intestinal barrier repair through multi-target integrated therapy.

At the engineering level, a BL composite system (INPs@BL@Gel) was constructed by integrating Bai, Tyr, and Inu with BL using a coordination assembly strategy. Bai and Tyr were incorporated through Fe(III) coordination to form carrier-free infinite coordination polymer nanoparticles (ICPs), enabling efficient loading of functional molecules and responsive release in inflammatory environments. The ICPs were subsequently coated with Inu and incorporated onto the BL surface to form a probiotic composite (INPs@BL). To enhance stability during oral delivery, the final system was encapsulated in a gelatin-silk fibroin hydrogel (SG-Gel), providing resistance to gastric acid and responsiveness to changes in pH and matrix metalloproteinase (MMP) activity. INPs@BL@Gel is designed for effective protection and targeted release in inflammation-associated sites within the gastrointestinal tract. This design not only realizes our proposed strategy but also takes full advantage of the multi-functional attributes of each component, such that no constituent is limited to a single function, ultimately yielding a structurally simplified yet functionally diversified INPs@BL@Gel.

In vivo and ex vivo experiments confirmed the exceptional biological activity and therapeutic efficacy of INPs@BL@Gel. Compared to the control group, its survival rate in gastric acid environments increased by approximately 56.87-fold, and its colonization in the inflamed intestine increased by about 10.58-fold, overcoming common challenges in oral probiotic delivery. INPs@BL@Gel significantly enhanced intestinal inflammation resolution, increasing colonic goblet cell abundance and restoring colon length to normal levels. It also modulated the expression of pro-inflammatory cytokines, such as TNF-α, IL-1β, and IL-6, while upregulating anti-inflammatory cytokine IL-10. Behavioral assessments revealed significant improvements in spontaneous activity, anxiety-like behavior, and reward responses in treated mice, with a reduction in immobility time in the tail suspension and forced swimming tests. Mechanistic investigations showed that the system reversed gut dysbiosis, restored SCFAs-producing bacteria, and corrected key metabolite imbalances, contributing to BBB repair and synaptic plasticity regulation. This study provides a new framework for oral, integrated intervention in colitis-associated depression and offers broader insights into probiotic engineering for gut-brain axis-related diseases.

## 2. Results and Discussions

### 2.1 Preparation and Characterization of INPs and INPs@BL@Gel

To increase the drug loading of the therapeutic drug and ensure its effective release at the site of enteritis, we chose to form ICPs by combining Bai, Tyr, and Fe^3+^. The infinite coordination polymer not only has a drug loading capacity close to 100% but also can be released at the inflammatory site. Subsequently, the polysaccharide Inu was coated on the surface. Since Inu is similar to Bai and easily combines with Bai, the intermolecular hydrogen bonds and hydrophobic interactions between them were utilized to deposit Inu π-π on the surface of ICPs nanoparticles. The above-mentioned infinite coordination and π-π stacking are both self-assembled carrier-free drug systems with large drug loading and controlled release at the inflammatory site, Finally, INPs@BL@Gel were obtained by ICPs nanoparticles coated on the bacterial system and then encapsulated in hydrogel formed by silk fibroin and gelatin (SG-Gel). SG-Gel has excellent acid resistance and has inflammatory site responsive release ability to meet the requirements for the multitargeting integrated therapy of intestinal inflammation.

As shown in Figure 1a, transmission electron microscopy (TEM) revealed that INPs exhibit a spherical morphology with an average particle size of 85 ± 11.5 nm. The TEM image clearly demonstrates the core-shell structure, where the dark core corresponds to the ICP structure, and the light-colored shell represents the Inu layer. Dynamic light scattering (DLS) measurements further confirmed the formation of INPs, with a hydrodynamic diameter of 113 ± 13.2 nm, which is larger than that of the ICPs alone (Figure 1b). The surface of the INPs also exhibited a positive charge (Figure 2b). To evaluate nanoparticle stability, INPs were stored phosphate buffer saline (PBS) for 60 days. During this period, their hydrated diameter remained consistently stable at approximately 118 nm (Supplementary Figure S1). No signs of aggregation or degradation were observed, indicating that INPs maintain excellent structural stability under physiological conditions. X-ray diffraction (XRD) analysis revealed a prominent diffraction peak at d = 3.4 Å, corresponding to the typical inter-aromatic ring distance involved in π-π stacking interactions, indicating that the self-assembly of INPs is driven primarily by π-π interactions (Figure 1e). UV-visible absorption spectra of Bai, Tyr, and their complexes with Fe^3+^ further confirmed the successful formation of the ion-conjugated ICPs. Following coordination, the absorption band of Bai at 224.51 nm shifted to 228.24 nm, while the characteristic peaks of Bai at 278.94 nm and Tyr at 322.65 nm were retained in the ICPs (Figure 1c). Infrared spectroscopy provided additional evidence of the coordination interaction. The C-H stretching vibration peak at 2899.12 cm^-1^ on the benzene ring showed enhanced intensity and a slight shift after ICP formation, indicating that hydrogen atoms on the aromatic ring are affected by the coordinating group during the coordination process. In the case of Tyr, the amino group (-NH[) band at 3203.70 cm^-1^ underwent a partial red shift after INP formation, further supporting the formation of coordination bonds (Figure 1d). X-ray photoelectron spectroscopy (XPS) analysis was performed to further substantiate these findings. Figure 1f shows that the nanoparticles contain C, O, and Fe. The Fe 2p binding energies of the nanoparticles (710.80 eV and 725.16 eV), as compared to an FeCl_3_ standard, were deconvoluted into four peaks at 709.10 eV, 711.58 eV, 721.90 eV, and 724.20 eV. The Fe 2p binding energies were lower than those of FeCl_3_, indicating that Fe(III) accepted lone-pair electrons from coordinating atoms, which is a hallmark of successful coordination. Additionally, the O 1s binding energies of the nanoparticles (531.9 eV and 531.2 eV) were higher than those observed for Bai alone (531.2 eV and 529.3 eV) and Tyr alone (531.5 eV and 529.6 eV), further confirming the successful formation of ICPs (Figure 1g). The cytotoxicity of the samples was evaluated in intestinal epithelial cells. Cell viability remained essentially unchanged across a broad range of INP concentrations, with only a minor proportion of cells exhibiting cell death at concentrations exceeding 1 mg mL^-1^. These findings indicate that INPs exhibit minimal cytotoxicity towards intestinal epithelial cells, causing only minor damage at high concentrations (Figure 1h). These results, along with the above physicochemical characterization, suggest that INPs possess favorable stability and biocompatibility for subsequent biological applications.

**Figure 1.**
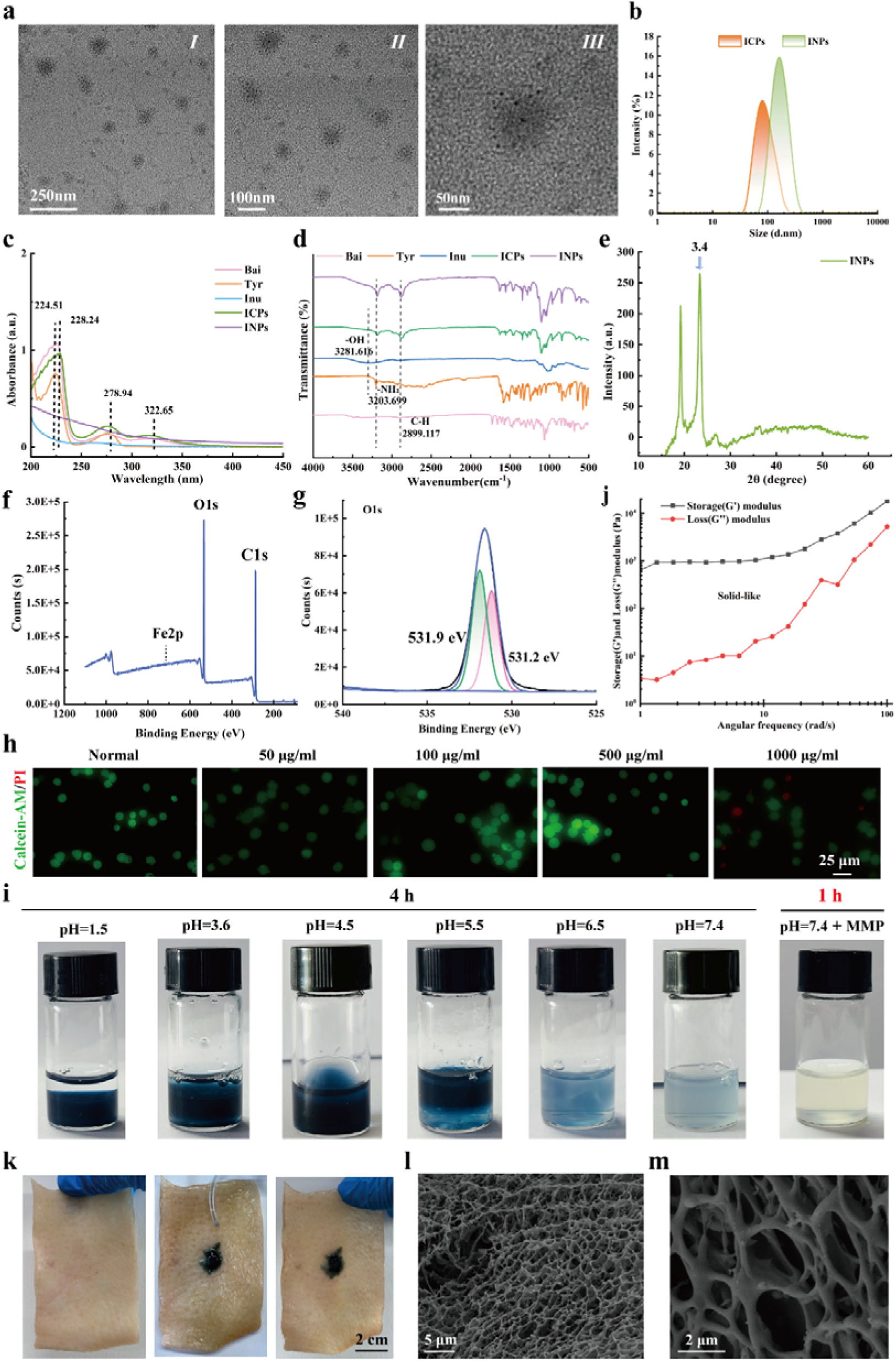
Characterization of INPs and SG-Gel. (a) Representative TEM images of INPs (with different magnifications). (b) Grain-size graph of ICPs and INPs. (c, d) Ultraviolet and Fourier infrared images of various compounds in INPs. (e) XRD result diagram of INPs. (f, g) The full spectrum graph of XPS in INPs and the peak-spectrum graph of the O element. (j) Rheological properties of SG-Gel. (h) The cytotoxicity of INPs. (i) The pH of SG-Gel and the response release duration of MMP. (k) The pigskin adhesion of SG-Gel. (l, m) Representative SEM images of SG-Gel (with different magnifications).

**Figure 2.**
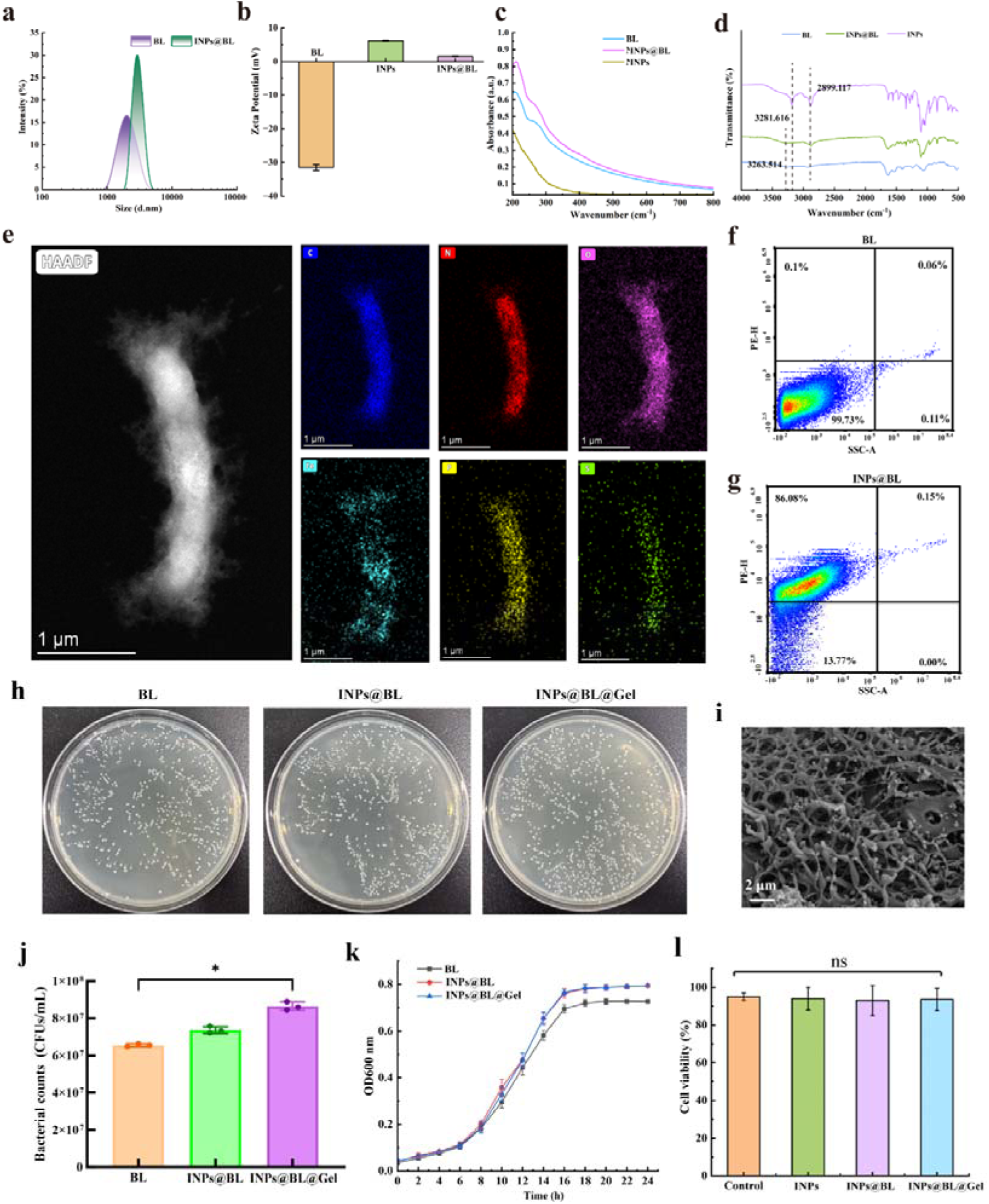
Characterization of INPs@BL@Gel. (a) The particle sizes of BL and INPs@BL. (b-d) The zeta potential, ultraviolet spectra and infrared spectra of BL, INPs and INPs@BL. (e) HAADF-STEM images and EDS spectra of INPs@BL. (f, g) Flow cytometry analysis of the encapsulation rate of INPs on BL. (i) Representative SEM image of INPs@BL@Gel. (h, j) Photographs and corresponding counts of bacterial colonies formed on TPY AGAR plates. (k)The proliferation ability of BL was evaluated by the OD600 values cultured in TPY medium at different time points, n=3. (l) Cell viability of NCM 460 cells treated with different drug groups (BL, INPs@BL and INPs@BL@Gel) for 2 hours. Data are expressed as Mean ± standard deviation (Mean ± SD), n=3. Statistical analysis was evaluated with two-tailed Student’s t tests (*p < 0.05, **p < 0.01, ***p < 0.001, and ****p < 0.0001).

For oral and intragastric administration, a hydrogel with suitable flow and gelation properties is essential for protection. To this end, we prepared a dark-blue SG-Gel by crosslinking silk fibroin extracted from silkworm cocoons with gelatin in the presence of genipin. Rheological measurements of SG-Gel revealed that the storage modulus (G’), reflecting elasticity, was consistently higher than the loss modulus (G”), which indicates viscosity, over the entire range of angular frequencies (Figure 1j). As angular frequency increased, G’ remained stable, with a slight increase, while G” gradually increased, suggesting that the material exhibited predominantly elastic behavior at lower frequencies, transitioning to a more viscous behavior at higher frequencies. The stability of SG-Gel under different pH conditions was further evaluated by incubating it in buffer solutions of varying pH values. As shown in Figure 1i and Supplementary Figure S2, under acidic conditions (pH 1.5), SG-Gel remained largely intact after 4 hours, with a residual gel weight of approximately 2.205 ± 0.05 g. As the pH increased, SG-Gel gradually dissolved, with gel weight decreases of 15.65%, 25.12%, 35.83%, and 78.59% at pH values of 3.6, 4.5, 5.5, and 6.5, respectively. At pH 7.4, the residual gel structure was barely visible, and the gel weight decreased to 0.024 ± 0.04 g, indicating that SG-Gel remains stable under strongly acidic conditions but undergoes gradual degradation as the environmental acidity weakens. Notably, in the presence of matrix metalloproteinase (MMP), SG-Gel completely degraded within one hour at pH 7.4, with its color changing from deep blue to transparent (Figure 1i). This enzyme-responsive behavior, which causes rapid disintegration in the presence of MMPs, highlights the exceptional environmental sensitivity of SG-Gel. Subsequently, the degradation behavior of SG-Gel was assessed in simulated gastric fluid (SGF) and simulated intestinal fluid (SIF). As shown in Supplementary Figure S3, SG-Gel remained largely intact in SGF, with minimal degradation observed over 72 hours. In contrast, SG-Gel gradually disintegrated in SIF, and its degradation rate in MMP-containing SIF was approximately four times faster than in pure SIF, demonstrating the dual responsiveness of SG-Gel to neutral pH and MMPs. This property enables SG-Gel to degrade in the intestinal environment, releasing its payload at the target site. The mucoadhesive properties of SG-Gel were further evaluated using porcine skin tissue. Figure 1k shows that when SG-Gel was applied to the epidermis of skin tissue, it adhered firmly without slipping, even when the tissue was placed in a vertical position for extended periods. These observations suggest that SG-Gel possesses strong mucoadhesive properties, facilitating prolonged retention in the intestine during intragastric administration. Finally, scanning electron microscopy (SEM) revealed that SG-Gel possesses a three-dimensional porous network structure, which is advantageous for loading various therapeutic agents (Figures 1l, m).

### 2.2 Characterization of INPs@BL@Gel

BL is a widely utilized probiotic known for its excellent biocompatibility and reported anti-inflammatory and microbiota-modulating effects, making it a promising candidate for the treatment of gastrointestinal disorders and associated neurobehavioral symptoms. Previous studies have demonstrated that BL can help restore the intestinal mucosal barrier and modulate immune responses. Additionally, its metabolites have been linked to neuroprotective effects and improved synaptic plasticity, which may contribute to alleviating depressive-like symptoms associated with colitis. Based on these findings, *B. longum ATCC BAA-999* was selected as the therapeutic strain, and INPs were incorporated onto its surface to enhance its therapeutic potential. INPs are rich in hydroxyl groups and amino acid structural units, which form the basis of their adhesive properties. These characteristics allow INPs to strongly adhere to surfaces such as cells, tissues, and mucosal membranes, facilitating the stable attachment of nanoparticles to bacterial envelopes. By encapsulating INPs on the surface of BL to form a multifunctional layer, the therapeutic potential of the bacteria was significantly enhanced, leading to the successful construction of the INPs@BL composite system.

As shown in Figure 2a, the hydrodynamic diameter of BL slightly increased after the loading of INPs, indicating successful surface modification of the bacteria. Zeta potential measurements confirmed that BL and INPs carry opposite charges, which highlights the importance of electrostatic interactions in their binding process (Figure 2b). UV-Vis spectroscopy provided additional evidence for the formation of INPs@BL. The spectrum of INPs@BL retained the characteristic absorption peak of BL at 275 nm and exhibited an additional absorption band at 205 nm, attributed to the synergistic contribution of both INPs and BL (Figure 2c). Fourier-transform infrared (FTIR) analysis further corroborated the successful assembly of INPs@BL. Specifically, the band at 3263.51 cm^-1^ in INPs@BL overlapped with that of BL, while the signal at 2899.12 cm^-1^ matched that of INPs, indicating the coexistence of spectral features from both components in the hybrid system (Figure 2d). Both silk fibroin (SF) and gelatin are proteins that absorb near the amide I (≈1650 cm^-1^), amide II (≈1540 cm^-1^), and amide III (≈1240 cm^-1^) bands. The position and intensity of these bands in SG-Gel (silk fibroin-gelatin hydrogel) differ from those of SF and gelatin alone, suggesting that the hydrogel contains structural characteristics of both proteins (Supplementary Figure S4). Additionally, broad peaks near ≈3300 cm^-1^ were observed, corresponding to O-H and N-H stretching vibrations (from proteins and water molecules). The wide peak in SG-Gel is indicative of the superimposed effect of both proteins. High-angle annular dark-field scanning transmission electron microscopy (HAADF-STEM) elemental mapping revealed a uniform distribution of Fe on the bacterial surface, confirming that the coordination-polymer nanomedicine layer had been successfully coated onto BL (Figure 2e). To further validate the presence of INPs on the bacterial surface, INPs were labeled with phycoerythrin (PE), and fluorescence intensity was quantified by flow cytometry. A clear PE signal was detected in INPs@BL, confirming the presence of INPs, with an estimated surface coverage of 85% (Figure 2f, g). INPs@BL was then encapsulated within the hydrogel to obtain INPs@BL@Gel. As shown in the scanning electron microscopy (SEM) cross-sectional image, the hydrogel matrix effectively embedded INPs@BL (Figure 2i). To confirm the viability of BL in INPs@BL and INPs@BL@Gel, bacterial suspensions were plated on agar after exposure to neutral pH and MMP to trigger gel disintegration. After 48 hours, the number of colonies did not decrease and even showed a slight increase, indicating that neither the INP coating nor the hydrogel encapsulation compromised bacterial viability (Figure 2h).

We next assessed whether INPs and the hydrogel affected BL proliferation. Figure 2k shows that the growth curves of INPs@BL and INPs@BL@Gel followed a trend comparable to that of unmodified BL over the same incubation period. Quantitative colony counting further supported this observation, with INPs@BL@Gel reaching 8.60 × 10^7^ CFU mL^-1^, INPs@BL reaching 7.35 × 10^7^ CFU mL^-1^, and BL reaching 6.55 × 10^7^ CFU mL^-1^, suggesting that surface coating and hydrogel encapsulation did not inhibit bacterial growth, and may have slightly promoted it (Figure 2j). Overall, the bacterial counts in the INPs@BL and INPs@BL@Gel groups were 1.12-fold and 1.31-fold higher than in the BL-only group, respectively. This modest increase may be due to BL’s ability to utilize Inu, as well as gelatin and silk fibroin in the hydrogel, as fermentable nutrient sources that support bacterial growth.

In simulated gastric fluid (SGF), the BL group decreased to approximately 4.22 × 10^5^ CFU mL^-1^ after 4 hours (Supplementary Figure S5a, b). In contrast, bacterial viability in SGF increased by 7.76-fold for INPs@BL and by 56.87-fold for INPs@BL@Gel, compared to BL. In simulated intestinal fluid (SIF), INPs@BL and INPs@BL@Gel demonstrated 1.59-fold and 10.58-fold higher viability than BL after 4 hours, respectively (Supplementary Figure S5c, d). These results highlight that SG-Gel significantly improves bacterial survival under gastric and intestinal conditions, facilitating the accumulation of INPs@BL at intestinal injury sites. Finally, cytotoxicity assays indicated that INPs, INPs@BL, and INPs@BL@Gel exhibit good biocompatibility (Figure 2l).

### 2.3 Intracellular and Extracellular Antioxidant Activities of INPs@BL@Gel

Inflammation in the colon and brain is often associated with elevated levels of intra- and extracellular oxidative stress. This exacerbates disease progression by disrupting the intestinal barrier and blood-brain barrier (BBB), promoting inflammatory responses, and inducing cell apoptosis. Therefore, reducing oxidative stress is essential for treating colitis-associated depression.

To evaluate the antioxidant capacity of INPs, we utilized UV-Vis spectrophotometry to assess their ability to scavenge various free radicals. As shown in Figure 3a and 3b, INPs effectively scavenged ABTS (2,2’-azobis (3-ethylbenzothiazoline-6-sulfonate)) and DPPH (1,1-diphenyl-2-picrylhydrazyl) free radicals in a concentration-dependent manner. As the concentration of INPs increased, the levels of free radicals gradually decreased. Consistent with these findings, the blue-green color of the ABTS solution faded to colorless upon mixing with INPs, with absorbance measured at 734 nm. Similarly, the purple DPPH solution gradually turned pale yellow, with absorbance measured at 519 nm.

**Figure 3.**
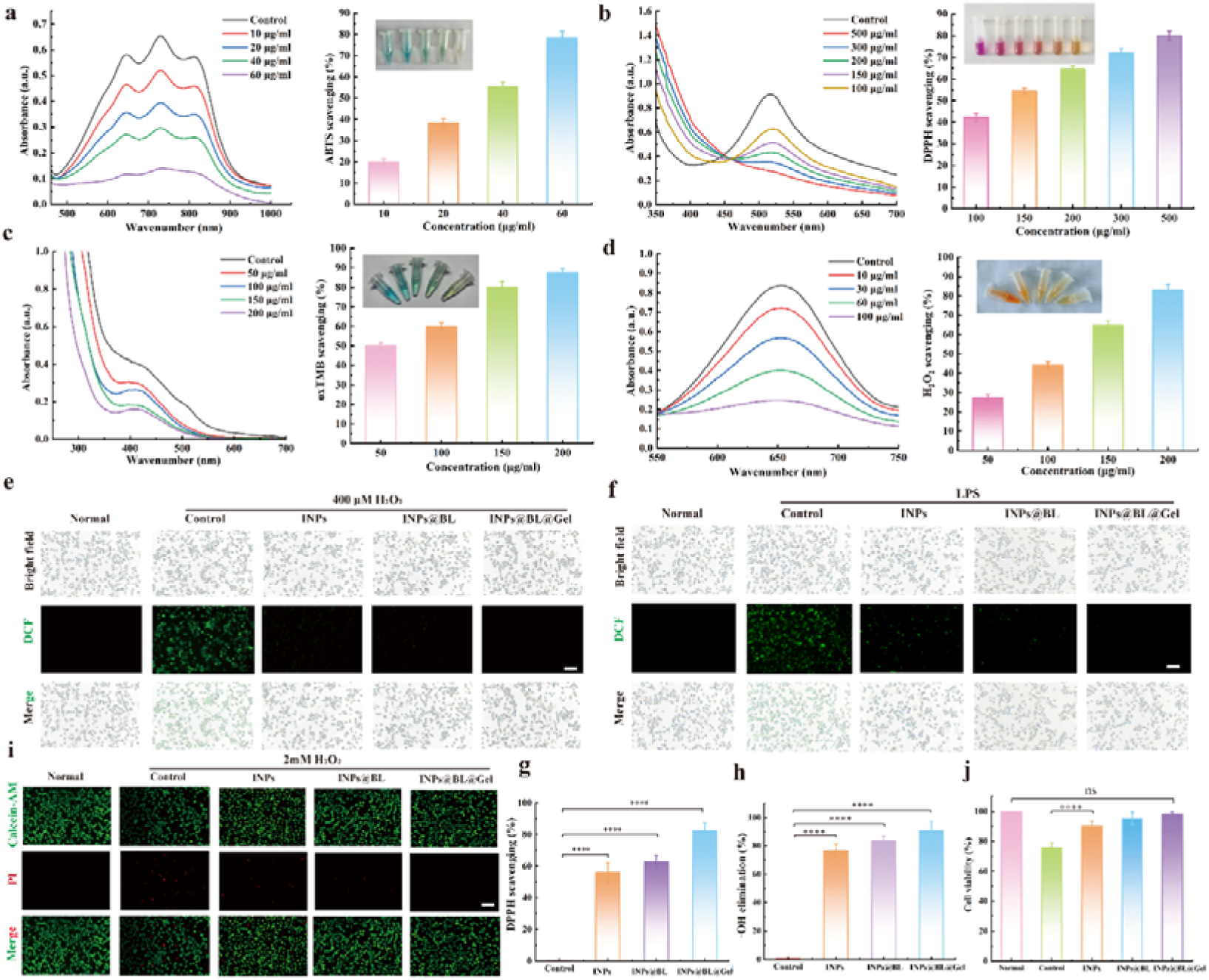
Determination of intracellular and extracellular antioxidant activity of INPs@BL@Gel. (a) The scavenging ability of ABTS free radicals at different INPs concentrations and the corresponding color changes. (b) The free radical scavenging ability of DPPH and the corresponding color changes at different INPs concentrations. (c) TMB was used to detect the scavenging ability of different INPs concentrations for •OH. (d) H_2_O_2_ was used to detect the scavenging ability of different INPs concentrations for O_2_^•—^. (e) The intracellular ROS level in NCM 460 cells was detected after exogenous H_2_O_2_ treatment under a fluorescence microscope. (f) The intracellular ROS level in NCM 460 cells was detected after endogenous LPS induction under a fluorescence microscope. (i) Calcein-AM/PI fluorescence images of NCM 460 cells treated with 2 mM H_2_O_2_ and various materials for 4 hours. Red fluorescence represents dead cells and green fluorescence represents live cells, and activity statistics (j). (g, h) In vitro clearance ability of DPPH and •OH by different material treatment groups. Data are expressed as means ± SD, n=3. Statistical analysis was evaluated with two-tailed Student’s t tests (*p < 0.05, **p < 0.01, ***p < 0.001, and ****p < 0.0001).

Furthermore, the total antioxidant activity of INPs under oxidative stress conditions was evaluated by TMB (3,3’,5,5’- Tetramethylbenzidine) and H_2_O_2_ removal assays. As shown in Figures 3c and 3d, as the concentration of INPs increased, the removal efficiency for TMB (color changing from blue to light yellow) and H_2_O_2_ (color changing from orange-red to pale yellow) gradually improved. These results demonstrate that INPs possess broad-spectrum antioxidant activity, capable of effectively scavenging reactive oxygen species (ROS) and mitigating cell damage caused by oxidative stress. Based on these findings, a concentration of 150 μg mL^-1^ was selected for subsequent experiments, with INPs@BL and INPs@BL@Gel prepared using equal INPs content.

Additionally, we employed 2’,7’-dichlorofluorescein diacetate (DCFH-DA) as a fluorescent probe to assess the intracellular ROS-scavenging activity of INPs, INPs@BL, and INPs@BL@Gel. As shown in Figures 3e and 3f, INPs significantly reduced the ROS levels in NCM 460 cells under both endogenous oxidative stress (induced by LPS) and exogenous oxidative stress (induced by H_2_O_2_). These results indicate that INPs effectively reduce the intracellular oxidative load and protect cells from oxidative damage. Notably, INPs@BL and INPs@BL@Gel exhibited stronger ROS-scavenging effects than INPs alone. This enhanced activity may be attributed to Bai, which can inhibit the formation of reactive oxygen/nitrogen species and neutralize free radicals through electron/hydrogen donation. By interrupting radical-driven chain reactions, Bai reduces the amplification of oxidative stress. Moreover, Bai can directly provide hydrogen atoms to active free radicals, further protecting cells from oxidative damage.

In vitro antioxidant experiments consistently demonstrated that INPs, INPs@BL, and INPs@BL@Gel (at the same drug dose) exhibited robust free radical scavenging activity against DPPH and hydroxyl free radicals (•OH) (Figures 3g, 3h). Among these, INPs@BL@Gel showed the highest clearing efficiency, with a clearing rate of 82.3% for DPPH and 91.0% for • OH. Live/dead staining further confirmed the protective effects of these formulations against oxidative injury. As shown in Figure 3i, INPs, INPs@BL, and INPs@BL@Gel treatment reduced H_2_O_2_-induced NCM 460 cell damage. Quantitative analysis revealed that 2 mM H_2_O_2_ reduced cell viability to 74.5%, while the viability of all treatment groups was maintained at over 90.0% (Figure 3j). Taken together, these findings demonstrate that INPs exhibit significant free radical scavenging and antioxidant activity. The synergy between BL and the hydrogel further enhances these effects, improving the body’s resistance to oxidative stress, reducing intracellular ROS production, and effectively protecting intestinal epithelial cells from oxidative damage.

### 2.4 Mucoadhesive Properties of INPs@BL@Gel

Stable and long-term intestinal colonization of probiotics is crucial for achieving sustained therapeutic benefits. The SG-Gel is designed to form a viscous network within the intestine, which adheres to the mucus layer, thereby strengthening contact with the intestinal wall and prolonging the residence time of the therapeutic payload at the target site. To evaluate the mucosal adhesion of INPs@BL@Gel in inflamed colon tissue, fluorescently labeled INPs were co-incubated with freshly excised intestinal segments from mice with colitis.

As shown in Figure 4a, the fluorescence intensity of the INPs@BL group increased compared to the INPs group, indicating that probiotics promoted particle retention within the intestine. Notably, the INPs@BL@Gel group exhibited significantly higher fluorescence than both the INPs and INPs@BL groups, demonstrating that the hydrogel further enhanced the adhesion of particles and BL to the intestinal mucosa. Fluorescence quantification confirmed this, revealing that the addition of SG-Gel effectively enhanced particle enrichment within the intestinal tissue.

**Figure 4.**
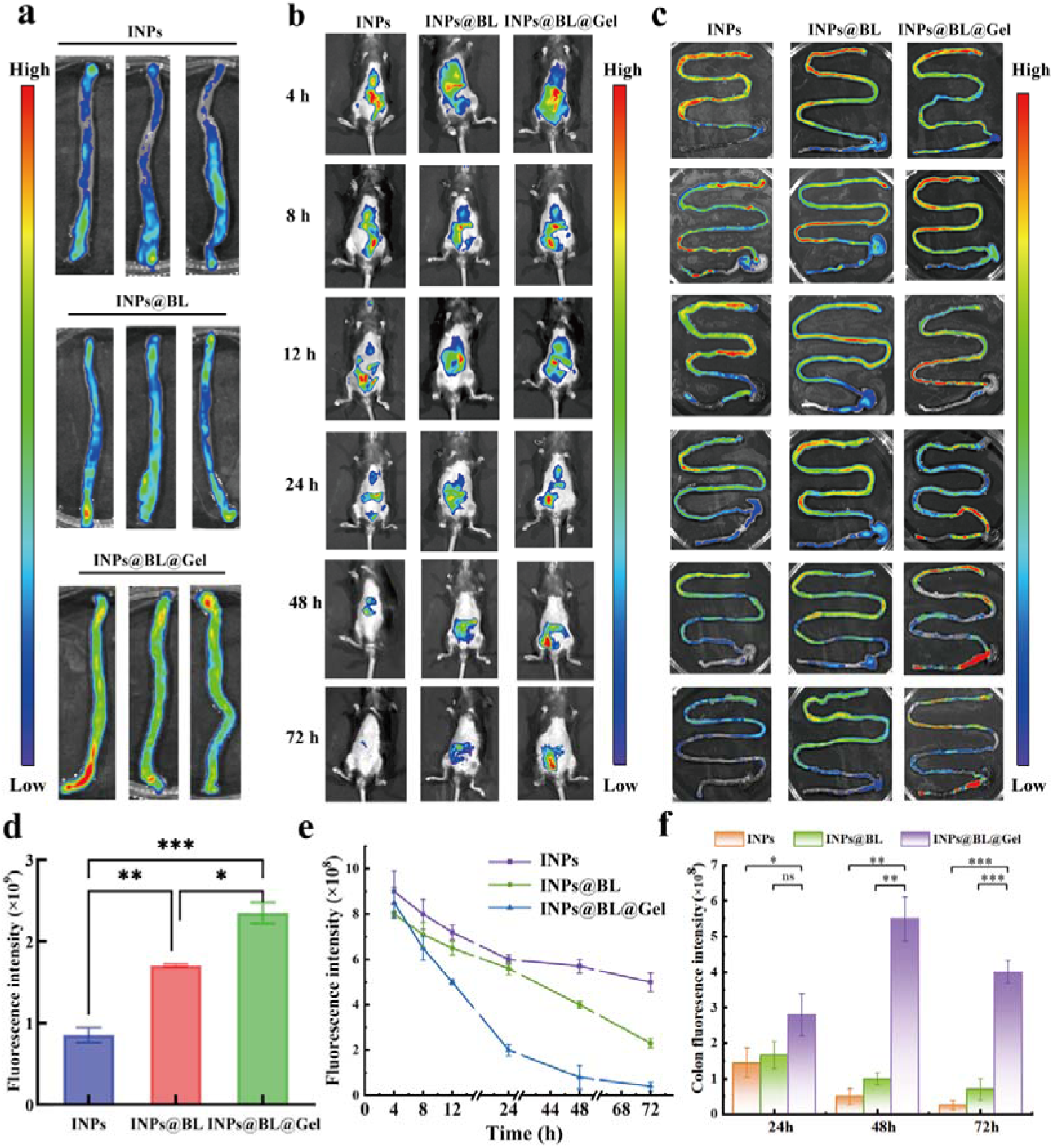
Mucosal adhesion ability of INPs@BL@Gel. (a) In vitro colonic fluorescence images after incubation with different drugs (INPs, INPs@BL and INPs@BL@Gel) marked with IR780. (b, c) Fluorescence imaging images of live mice and intestinal tracts in vivo treated with different drug groups at different time points. (d) Fluorescence intensity statistics of the colon in vitro. (e) Fluorescence intensity attenuation curve in mice. (f) Fluorescence intensity of the colon of mice at different time points. Data are expressed as Mean ± SD, with n=3. Statistical analysis was evaluated with two-tailed Student’s t tests (*p < 0.05, **p < 0.01, ***p < 0.001, and ****p < 0.0001).

To observe the distribution of the materials more precisely, mice were euthanized at specific time points, and their intestines were isolated for fluorescence imaging. As shown in Figure 4f, at 48 hours, the fluorescence intensity in the INPs@BL@Gel group was 11.02 and 5.53 times greater than in the INPs and INPs@BL groups, respectively. By 72 hours, the fluorescence intensity in the INPs@BL@Gel group was 15.38-fold and 5.71-fold higher than in the INPs and INPs@BL groups, respectively. These results collectively demonstrate that incorporating BL and SG-Gel significantly prolongs and maintains drug retention within the gastrointestinal tract.

Further fluorescence imaging was performed to assess the material distribution within visceral organs. The results revealed no detectable fluorescence signals in major organs such as the heart, liver, spleen, lungs, and kidneys (Supplementary Figure S6). This suggests that the material undergoes primary metabolism within the gastrointestinal tract following oral administration, with minimal diffusion to visceral organs. Notably, the fluorescence intensity in the INPs@BL and INPs@BL@Gel groups increased to 2.13-fold and 2.65-fold, respectively, compared to the INPs group (Figure 4d). In vivo fluorescence imaging in mice revealed that fluorescence intensity gradually decreased over time in all groups (Figure 4b, c). However, the INPs@BL@Gel group exhibited significantly higher fluorescence intensity at each time point compared to the INPs and INPs@BL groups. Furthermore, the INPs@BL@Gel group displayed a markedly slower decline in fluorescence intensity than the other groups, maintaining strong fluorescence even after 72 hours. Quantitative fluorescence decay curves showed that the INPs group experienced a rapid decline in fluorescence intensity within 24 hours. In contrast, the INPs@BL and INPs@BL@Gel groups exhibited slower decay rates. These findings suggest that BL and SG-Gel effectively prolong the retention time of particles within the intestine. Notably, the INPs@BL@Gel group exhibited the highest fluorescence intensity at every time point and the slowest decay rate. This can likely be attributed to BL’s role as an intestinal colonizing microbiota that adheres strongly to the intestinal mucosal layer. Additionally, the targeted release of the hydrogel at sites of intestinal inflammation further enhances its therapeutic efficacy.

### 2.5 Therapeutic Efficacy of INPs@BL@Gel against DSS-Induced Colitis

To evaluate the in vivo therapeutic potential of INPs@BL@Gel, we conducted a comprehensive assessment in a mouse model of colitis induced by 3% dextran sulfate sodium (DSS). Mice were randomly assigned to five groups and, starting on day 7 after DSS administration, received daily treatments with PBS, INPs, INPs@BL, or INPs@BL@Gel for 5 consecutive days (Figure 5a). A healthy control group was included and gavaged with PBS for the same duration. After the treatment period, mice were euthanized, and their tissues were collected for subsequent analyses.

**Figure 5.**
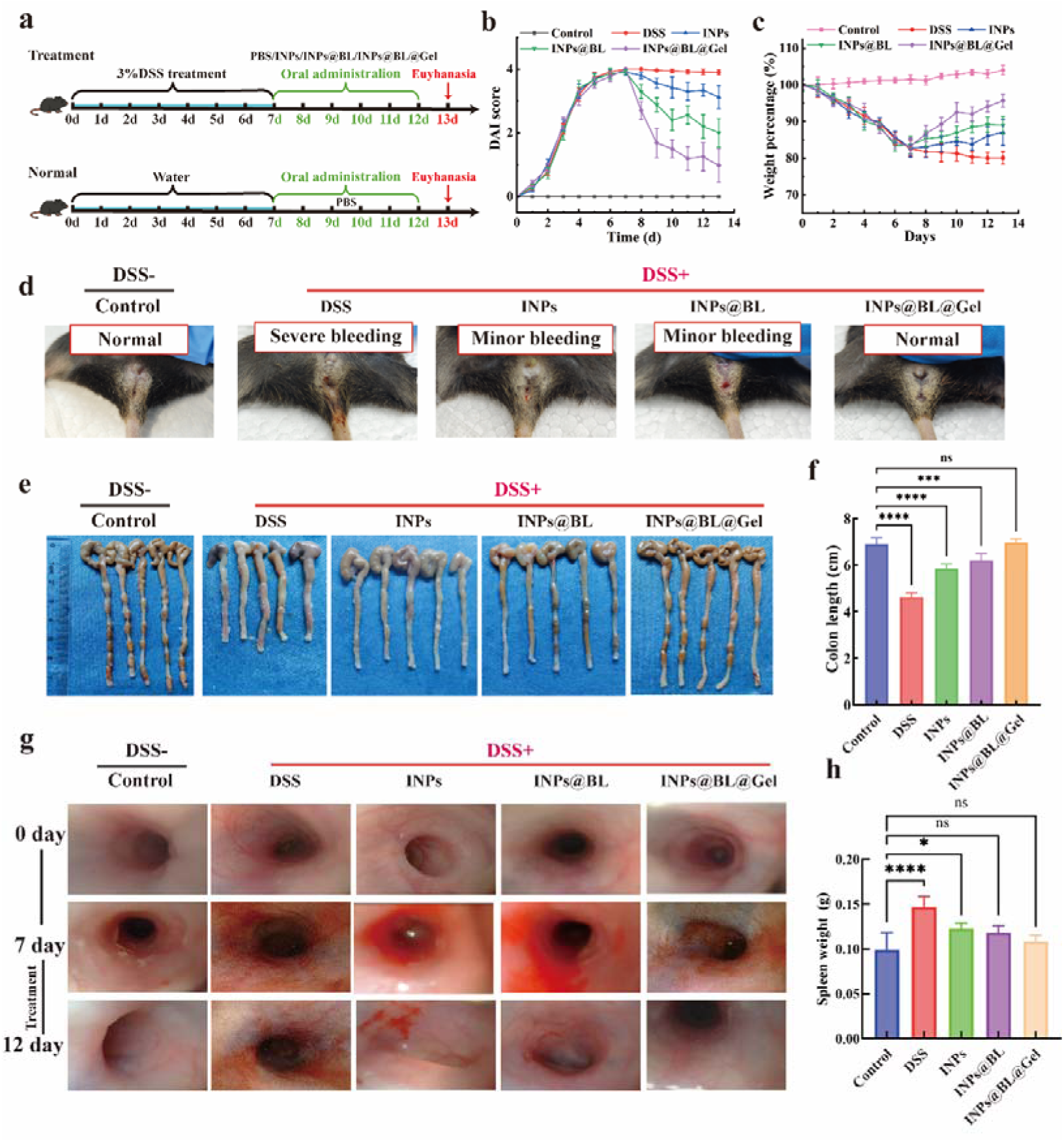
The treatment effect of INPs@BL@Gel on colitis. (a) Schematic diagram of modeling and treatment process. (b) Changes in the DAI of mice treated with different drug groups. (c) Changes in the body weight score of mice during the experiment. (d) Representative photos of the anus of each group of mice. (e) Photographs of the colons and (f) the statistics of colon lengths for each group of mice. (g) Endoscopic images of the colon in each group of mice. (h) Quantitative analysis of spleen weight in different groups of mice. Data are expressed as Mean ± SD, with n=5. Statistical analysis was evaluated with two-tailed Student’s t tests (*p < 0.05, **p < 0.01, ***p < 0.001, and ****p < 0.0001).

Throughout the study, body weight, colonic condition, and the disease activity index (DAI)—which includes measurements of weight loss, stool consistency, and rectal bleeding—were carefully monitored. As shown in Figure 5b, 5c, and Supplementary Table S2, DSS-induced mice exhibited a significant loss of body weight and a marked increase in the DAI, indicating severe colitis. In contrast, all treatment groups demonstrated varying degrees of colitis alleviation. Notably, mice treated with INPs@BL@Gel exhibited a clear rebound in body weight during the later stages of treatment, accompanied by a significant reduction in the DAI, approaching the levels observed in healthy controls. Additionally, DSS exposure induced prominent clinical symptoms, including rectal bleeding and diarrhea, whereas INPs@BL@Gel treatment alleviated these symptoms to the greatest extent (Figure 5d). On day 13, the mice were euthanized, and colonic tissues were collected for further analyses. Colon shortening is a well-established indicator of colonic inflammation (Figure 5e). Ex vivo measurements revealed that compared to the control group (6.90 cm), colon length was reduced by 33.04%, 15.36%, and 10.14% in the DSS (4.62 cm), INPs (5.84 cm), and INPs@BL groups (6.20 cm), respectively. However, the reduction in the INPs@BL@Gel group (6.96 cm) was not significantly different from that of healthy controls (Figure 5f). Spleen weight analysis revealed marked splenomegaly in DSS-induced mice, with spleen weight approximately 1.50-fold higher than that of healthy mice. In contrast, no significant spleen enlargement was observed in the INPs@BL@Gel group, which was comparable to the control group (Figure 5h and Supplementary Figure S7). In addition, colonoscopy was performed to monitor mucosal changes throughout the treatment period. As shown in Figure 5g, DSS-induced mice exhibited severe inflammation, including mucosal bleeding, distorted vascular patterns, mucosal opacity, and adhesions. Notably, INPs@BL@Gel treatment led to a substantial normalization of the mucosal appearance, with colonoscopic findings comparable to healthy controls. In comparison to a previous report by the Sung-Kwon Moon team, which showed a 19.30% recovery in colon length after treatment (reaching approximately 4.6 cm), our treatment group demonstrated a recovery to approximately 6.0 cm, a value that was not significantly different from the normal group.^[39]^ This outcome highlights the remarkable therapeutic effect of INPs@BL@Gel.

Collectively, in the DSS-induced colitis model, INPs, INPs@BL, and INPs@BL@Gel all demonstrated therapeutic benefits to varying degrees, with INPs@BL@Gel consistently achieving the most pronounced improvement. Across multiple readouts, the INPs@BL@Gel group approached the levels observed in healthy controls, suggesting a synergistic therapeutic effect among the components and validating the feasibility of our design strategy.

### 2.6 Colonic Barrier Restoration and Inflammation Resolution in DSS-Induced Colitis

The loss of mucin-producing goblet cells and disruption of the mucus barrier are key events in the onset and progression of UC, contributing significantly to disease exacerbation. To evaluate intestinal barrier integrity and the mechanisms underlying barrier damage, we analyzed intestinal tissue and blood samples from the DSS, INPs, INPs@BL, INPs@BL@Gel, and healthy control groups.

Histological evaluation was first performed using Hematoxylin and Eosin (H&E) staining, with tissue damage quantified via a comprehensive histological scoring system that considers inflammation severity, lesion depth, crypt damage, and the extent of the affected area (Supplementary Table S3). As shown in Figure 6a, DSS-induced colitis led to severe colon damage, characterized by significant lymphocyte infiltration and crypt destruction, with a histological score of approximately 9. In contrast, all treatment groups showed significant tissue repair, with the INPs@BL@Gel group demonstrating the most remarkable restoration of epithelial integrity, with a histological score reduced to about 1 point (Figure 6b).

**Figure 6.**
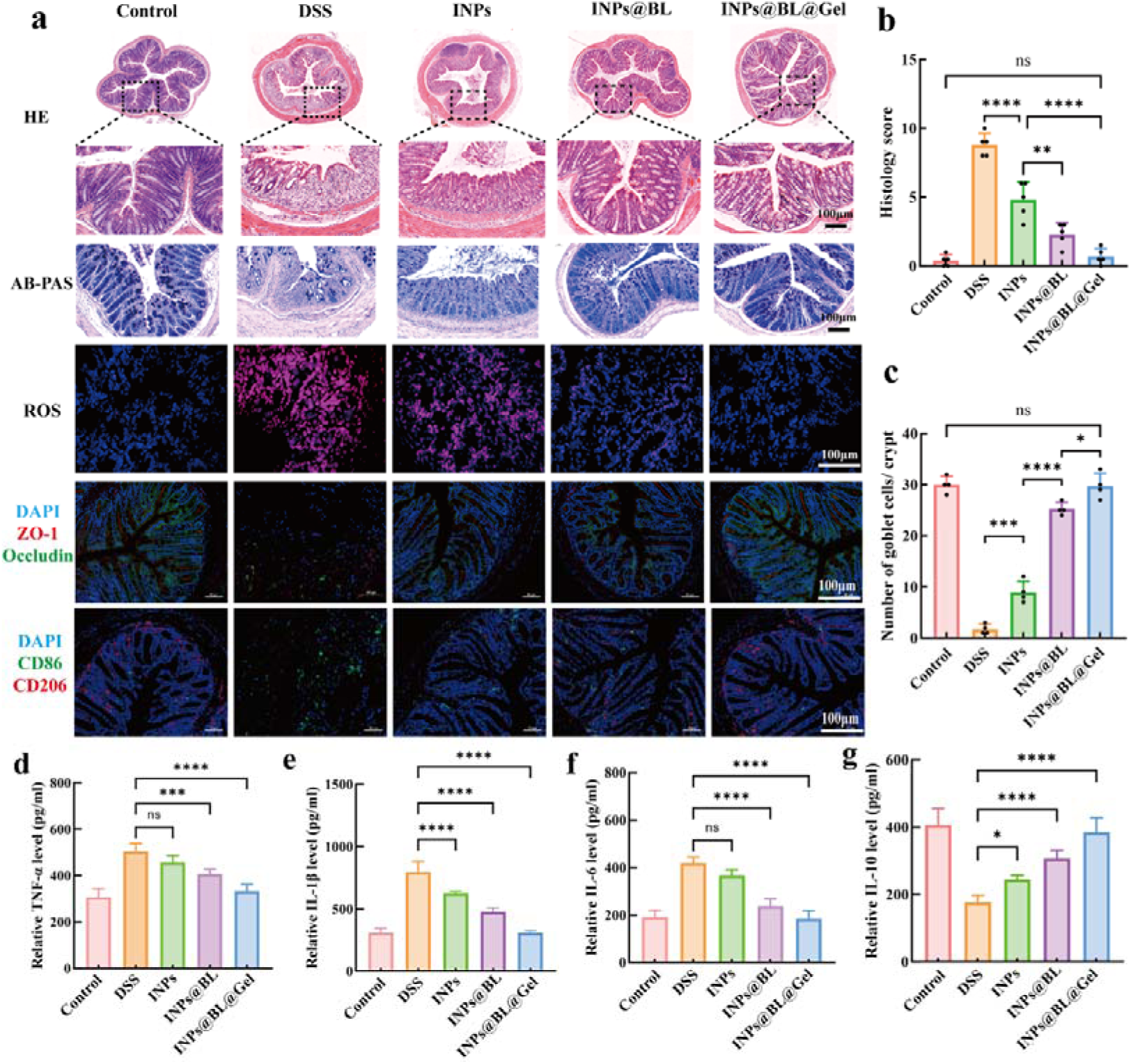
INPs@BL@Gel for the repair of colonic barrier damage and the clearance of inflammation. (a) Representative H&E, AB-PAS and ROS staining images of the colon tissue of mice treated with different drug groups; And the immunofluorescence co-localization images of tight junction proteins (ZO-1, Occludin) and macrophage markers (CD86, CD206) in the colon. (b) Histological injury scores of colon tissue after treatment with different drug groups. (c) The number of goblet cells in the colonic tissues of mice in each group. (d-g) The levels of pro-inflammatory cytokines (TNF-α, IL-1β and IL-6) and anti-inflammatory cytokines (IL-10) in the colon tissue of mice in each group were determined by ELISA. Data are expressed as Mean ± SD, with n = 5. Statistical analysis was evaluated with two-tailed Student’s t tests (*p < 0.05, **p < 0.01, ***p < 0.001, and ****p < 0.0001).

Subsequently, we assessed the ability of INPs@BL@Gel to repair the intestinal mucosal barrier. Alcian blue-periodic acid-Schiff (AB-PAS) staining showed that INPs@BL@Gel effectively reversed the inflammation-induced damage to the mucus layer, restoring it to a normal state. This restoration was accompanied by a marked increase in the number of goblet cells and the thickening of the mucus layer (Figure 6a, Supplementary Figure S8). Compared with the DSS-induced colitis group, the number of goblet cells in the INPs@BL@Gel group increased approximately 20-fold (Figure 6c). To quantify intestinal reactive oxygen species (ROS) levels, we used fluorescent imaging based on dihydroethidium (DHE). In all treatment groups, INPs@BL@Gel exhibited the lowest fluorescence intensity, similar to the healthy control group, indicating its strong ROS scavenging ability (Figure 6a, Supplementary Figure S9). Furthermore, INPs@BL@Gel significantly increased the expression of tight junction proteins ZO-1 and Occludin, further supporting the repair of the intestinal barrier (Figure 6a, Supplementary Figure S10).

We also explored the potential anti-inflammatory effects of INPs@BL@Gel in DSS-induced intestinal inflammation. Macrophages, as key immune cells, play a crucial role in shaping inflammatory responses. Thus, studying the effect of INPs@BL@Gel on macrophage behavior may help elucidate its treatment mechanism in intestinal inflammation. Promoting the phenotypic shift from pro-inflammatory M1 macrophages to anti-inflammatory M2 macrophages is considered a promising strategy to restore immune homeostasis in inflammatory bowel diseases (IBDs). CD86 is a surface marker for M1 macrophages, while CD206 is widely used to identify M2 macrophages. Immunofluorescence staining of colon sections showed that CD86 expression was extremely low in healthy mice, indicating immune homeostasis (Figure 6a, Supplementary Figure S11). In contrast, DSS stimulation led to the dominance of CD86 signals in colon tissue, suggesting that macrophages shifted to the pro-inflammatory M1 phenotype. In contrast, the treatment groups showed a reduction in CD86 expression, accompanied by enhanced CD206 staining. Notably, the INPs@BL@Gel group showed negligible CD86 signals, consistent with immune homeostasis recovery and a tissue microenvironment conducive to repair.

To further verify the anti-inflammatory mechanism, enzyme-linked immunosorbent assay (ELISA) was used to detect levels of representative pro-inflammatory and anti-inflammatory cytokines in colon tissue, including TNF-α, IL-1β, IL-6 (pro-inflammatory), and IL-10 (anti-inflammatory). As shown in Figures 6d-f, DSS challenge significantly elevated the levels of TNF-α, IL-1β, and IL-6 to 1.64-, 2.57-, and 2.18-fold those in healthy controls, respectively. Conversely, the anti-inflammatory cytokine IL-10 was reduced by 56.83% compared to healthy mice, consistent with the heightened inflammatory state in colitis (Figure 6g). Following treatment with INPs, INPs@BL, or INPs@BL@Gel, colonic inflammation was markedly alleviated: the pro-inflammatory cytokines TNF-α, IL-1β, and IL-6 were significantly suppressed, while IL-10 levels were increased. Notably, compared with the DSS group, INPs@BL@Gel reduced TNF-α, IL-1β, and IL-6 levels in colonic tissue by 33.90%, 61.31%, and 55.82%, respectively, and increased IL-10 by 54.48%, outperforming both INPs and INPs@BL. These results are consistent with findings from the Yanling Hao team, who reported that *Bifidobacterium lactic acid* intervention reduced pro-inflammatory cytokines TNF-α, IL-1β, and IL-6 by 48.6%, 45.5%, and 24.5%, respectively.^[40]^ In comparison, the anti-inflammatory effect of our combination strategy with probiotics is more pronounced.

Collectively, these results demonstrate that INPs@BL@Gel not only exhibits significant therapeutic effects in DSS-induced colitis but also promotes barrier repair and inflammation resolution, achieving the best mucosal repair effect while also modulating immune responses and suppressing inflammation.

### 2.7 Alleviation of Colitis-Associated depression

DSS-induced colitis mice often exhibit significant anxiety- and depression-like behaviors, such as reduced motor activity and weakened exploration. This section evaluates whether oral INPs@BL@Gel can alleviate anxiety- and depression-like behaviors associated with colitis through five behavioral paradigms: the open field test (OFT), elevated plus maze (EPM), sucrose preference test (SPT), tail suspension test (TST), and forced swimming test (FST). Mice were given 3% DSS to induce colitis for 7 days, followed by oral treatment with PBS, INPs, INPs@BL, or INPs@BL@Gel for 5 consecutive days. The anxiety- and depression-like behaviors were then assessed through behavioral tests (Figure 7a).

**Figure 7.**
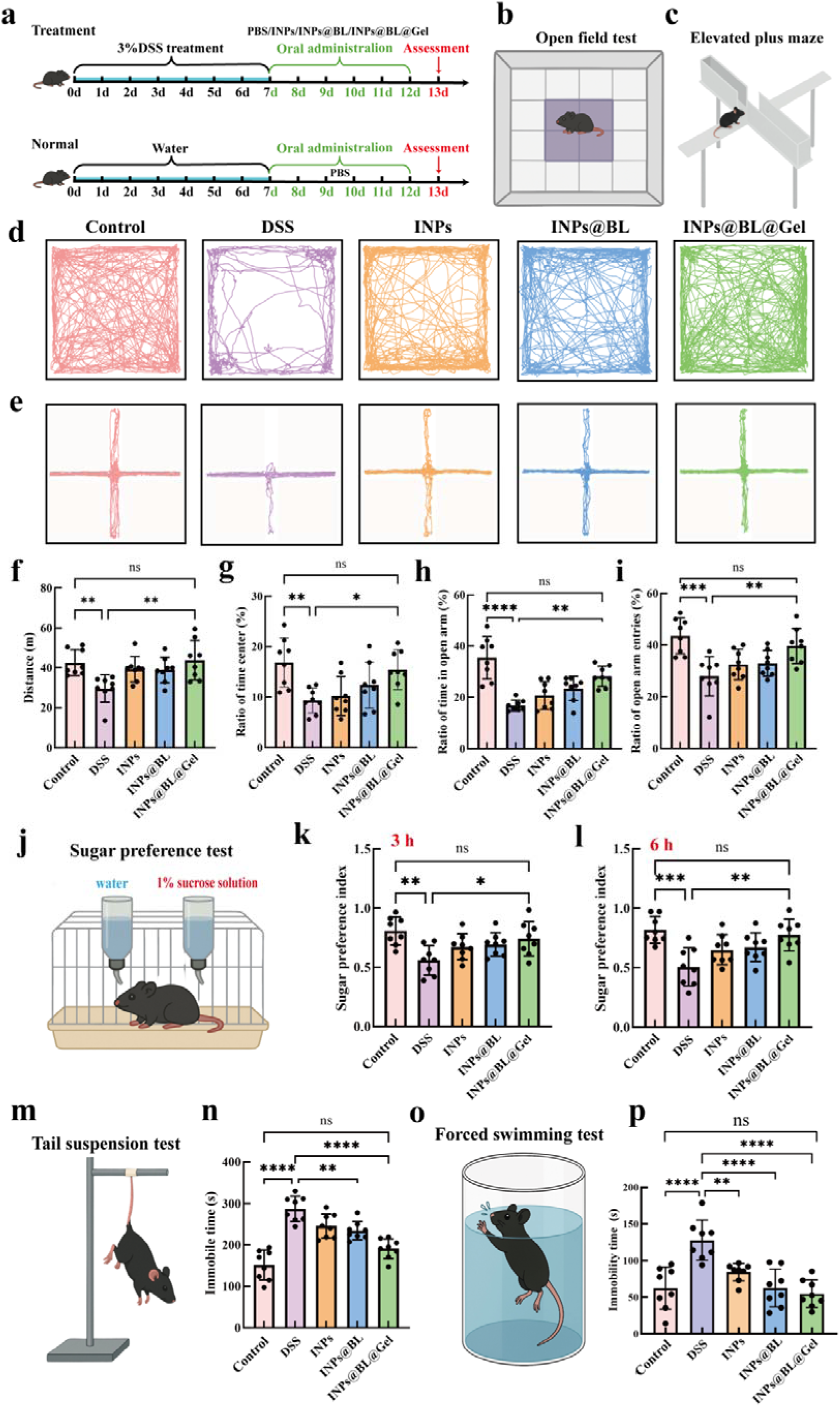
Behavioral Analysis of INPs@BL@Gel in Treating colitis-associated depression. (a) Schematic diagram of modeling and treatment process. (b, c) OFT schematic diagram and EPM schematic diagram.(d) Representative images of motion trajectory tracking of mice treated with different drug groups in OFT. (e) Representative images of the movement trajectory tracking of mice in each group in EPM. (f) Total movement distance of mice in each group of OFT. (g) Proportion of central time of mice in each group of OFT. (h) The proportion of time that mice in each group entered the arm-opening stage in EPM. (i) The proportion of the number of times mice in each group entered the open arm in the EPM. (j) Schematic diagram of SPT. (k, 1) The sugar water preference rates of mice in each group at 3 hours and 6 hours in SPT. (m,n) TST schematic diagram and the stationary time of each group. (o, p) FST schematic diagram and the stationary time of each group. Data are expressed as Mean ± SD, with n=8. Statistical analysis was evaluated with two-tailed Student’s t tests (*p < 0.05, **p <0.01, ***p < 0.001, and ****p < 0.0001).

In the OFT, DSS-induced mice exhibited stereotyped movement patterns, rarely entering the central area and moving short distances (Figure 7b, d). Quantitative analysis revealed that the DSS group’s total movement distance was reduced by nearly one-third, and their proportion of time spent in the central area was halved compared to the healthy control group (Figure 7f, g). After treatment, mice in the INPs, INPs@BL, and INPs@BL@Gel groups displayed improved exploratory behavior and reduced repetitive trajectories. Notably, the INPs@BL@Gel group showed the most significant improvement, with a total distance of ∼43.83 m and a central time ratio of 15.45%, approaching healthy control levels, indicating significant reductions in anxiety- and depression-like behaviors (Figure 7d).

In the EPM test, DSS-induced mice showed reduced time spent on the open arms and fewer open arm entries, indicating heightened anxiety. Mice treated with INPs@BL@Gel exhibited the strongest anti-anxiety effect, with a 1.68-fold increase in open arm stay time and a 1.35-fold increase in the number of open arm entries compared to the DSS group (Figure 7c, e, h, i).

In the SPT, the amount of sucrose consumed during the first 3 hours indicated the immediate reward response to initial exposure. As shown in Figure 7j, k, the DSS group’s preference for sucrose was about one-third lower than that of the healthy control group, reflecting a weakened interest in rewarding stimuli. After 6 hours, this preference decreased further by approximately two-fifths, indicating a persistent anhedonic-like behavior. Remarkably, INPs@BL@Gel treatment largely normalized this behavior, with sucrose preference indices of 0.74 and 0.78 at 3 hours and 6 hours, respectively, approaching the levels of healthy mice. These results suggest that DSS-induced mice developed a pronounced anhedonia-like phenotype, a core feature of depression, which was significantly improved by all treatment groups.

In the TST and FST, DSS-induced mice exhibited significantly prolonged immobility times: TST immobility was approximately 1.93-fold and FST immobility was 2.17-fold greater than that of healthy controls. Notably, INPs@BL@Gel treatment reduced immobility by 37.9% in the TST and 62.96% in the FST, showing a more pronounced effect than the other treatment groups (Figure 7m-p). The Honghong Yao team previously reported a reduction in immobility time by approximately one-third in NLRP3-deficient mice after microbiota treatment.^[41]^ In contrast, our treatment group exhibited a 37.9% and 62.96% reduction in TST and FST immobility times, respectively, highlighting the significant therapeutic effect of INPs@BL@Gel in alleviating anxiety- and depression-like behaviors.

Taken together, these behavioral assessments consistently demonstrate that DSS exposure induces anxiety- and depression-like behaviors in mice. INPs@BL@Gel significantly alleviates these colitis-associated depression. These findings further support the role of the gut-brain axis in mediating the therapeutic effects of INPs@BL@Gel on colitis-associated depression.

### 2.8 Restoration of BBB Integrity and Neural Function

Based on the behavioral outcomes, we hypothesized that DSS-induced colitis-associated depression may result from the leakage of gut lumen-derived molecules (GLDMs) and toxins from inflamed colonic tissue into the circulation. These circulating factors could initially impair blood-brain barrier (BBB) integrity, leading to neuroinflammation and local immune responses in the brain. To investigate this hypothesis, we evaluated BBB integrity using transmission electron microscopy (TEM) and Evans Blue (EB) extravasation experiments. As shown in Figure 8a, TEM images revealed significant ultramicrostructural abnormalities in the BBB of DSS-induced mice, including basal membrane destruction, disruption of tight junctions, and swelling of astrocytic glial cells. EB staining further confirmed the increased accumulation of EB in the brain, indicating enhanced BBB permeability (Supplementary Figure S12). In contrast, oral INPs@BL@Gel effectively reversed these pathological changes and maintained BBB structural integrity.

**Figure 8.**
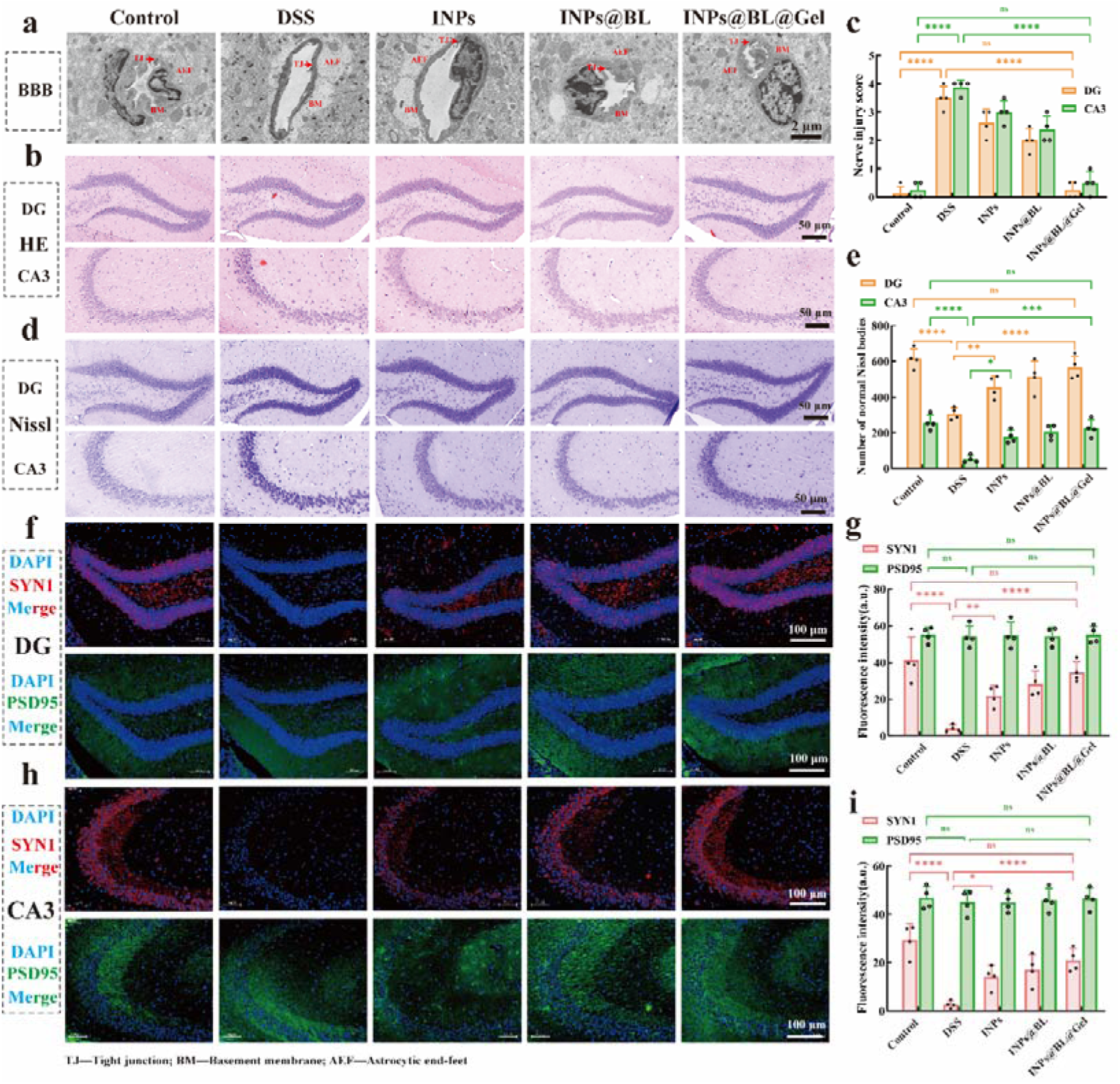
INPs@BL@Gel promotes neuronal repair and synaptic plasticity. (a) Representative TEM images of the BBB in mice after treatment with different drug groups. (b) Representative H&E staining images and nerve injury scores of the DG and CA3 regions in the hippocampus of mice after treatment with different drug groups (c). (d) Representative Nissl staining of active neurons in the DG and CA3 regions of the hippocampus in each group of mice and the number of intact Nissl bodies (e). (f) Representative immunofluorescence co-localization images and quantitative analysis of synaptic associated proteins (SYN1 and PSD95) in the DG region of the hippocampus of mice in each group (g). (h) Representative immunofluorescence co-localization images and quantitative analysis of synaptic associated proteins (SYN1 and PSD95) in the hippocampal CA3 region of mice in each group (i). Data are expressed as Mean ± SD, with n = 4. Statistical analysis was evaluated with two-tailed Student’s t tests (*p < 0.05, **p < 0.01, ***p < 0.001, and ****p < 0.0001).

H&E staining, along with Nissl staining, showed significant damage to the hippocampal region in DSS-induced mice (Figures 8b, d). The dentate gyrus (DG) and CA3 regions exhibited a decrease in the number of surviving neurons, neuronal degeneration, reduced Nissl bodies, and lower neuronal density compared to healthy controls. However, no obvious pathological alterations were observed in the DG or CA3 regions of INPs@BL@Gel-treated mice. Quantitative analyses of the H&E and Nissl staining results corroborated these findings (Figure 8c, e, Supplementary Table S4). The histological injury scores in the DSS group were approximately 3.5 (DG) and 3.875 (CA3). After INPs@BL@Gel treatment, these scores decreased to 0.25 (DG) and 0.5 (CA3), approaching those observed in healthy controls. Moreover, the number of intact Nissl bodies in the DG and CA3 regions of DSS-induced mice decreased to approximately half and one-fifth of that in healthy controls, respectively, while INPs@BL@Gel treatment increased these values by 1.837-fold (DG) and 4.28-fold (CA3) compared to the DSS group. These results indicate that INPs@BL@Gel treatment significantly reverses neuronal damage and structural alterations in the hippocampus caused by DSS-induced colitis.

To further assess synaptic plasticity, we performed immunofluorescence staining to examine the expression of the presynaptic protein synapsin-1 (SYN1) and the postsynaptic density protein PSD95 in the hippocampal DG and CA3 regions. Changes in pre- and postsynaptic markers provide insights into synaptic remodeling. As shown in Figures 8f, h (and Supplementary Figures S13 and S14), DSS exposure markedly reduced SYN1 expression in both the DG and CA3 regions. Normalized fluorescence intensity analysis revealed that SYN1 fluorescence in the CA3 region decreased by 90.65% in the DSS group compared to controls. Relative to the DSS group, SYN1 fluorescence increased by 5.14-fold (INPs), 6.62-fold (INPs@BL), and 7.64-fold (INPs@BL@Gel) (Figure 8i). A similar trend was observed in the DG region: compared to controls, SYN1 fluorescence in the PBS-treated (DSS) group decreased by 98.39%, while INPs, INPs@BL, and INPs@BL@Gel treatments increased SYN1 fluorescence by 5.36-fold, 6.95-fold, and 8.59-fold, respectively (Figure 8g, Supplementary Figure S13, S14). Notably, PSD95 staining showed weak expression in both DG and CA3 regions across all groups, with no significant differences in fluorescence intensity. Additionally, high-performance liquid chromatography (HPLC) analysis of INPs@BL@Gel revealed a well-resolved, baseline-separated peak at approximately tR ≈ 12.2 min, matching the characteristic peak of the HVA standard (Supplementary Figure S15). This suggests that INPs are metabolized by BL to produce HVA, which may contribute to preserving presynaptic integrity in the hippocampus and alleviating synaptic dysfunction associated with anxiety- and depression-like states.

In summary, our findings provide mechanistic insights into DSS-induced colitis-associated depression and demonstrate that INPs@BL@Gel is an effective strategy to preserve BBB integrity, improve synaptic plasticity, and potentially restore neural function.

### 2.9 Metagenomic Analysis

The gut microbiota plays a critical role in the pathogenesis of colitis-associated depression. Dysbiosis in this context is typically characterized by reduced microbial diversity and the expansion of pathogenic or opportunistic symbionts. To investigate the impact of the engineered INPs@BL@Gel on gut microbial composition, metagenomic sequencing was performed. Principal coordinates analysis (PCoA) based on species-level abundance profiles revealed clear differences in microbial community structure among groups (Figure 9a). The DSS-treated group was distinctly separated from the healthy control group, indicating severe dysbiosis induced by colitis. In contrast, INPs@BL@Gel treatment markedly reversed this dysregulation, with samples clustering closer to those of the control group in the PCoA space, suggesting a substantial recovery of the gut microbiota toward a homeostatic state. Phylogenetic analysis (Figure 9b) combined with a Kruskal-Wallis test (Figure 9c) further identified eight discriminant species that contributed most strongly to the observed intergroup differences, supporting partial but targeted restoration of the microbial ecosystem following INPs@BL@Gel administration.

**Figure 9.**
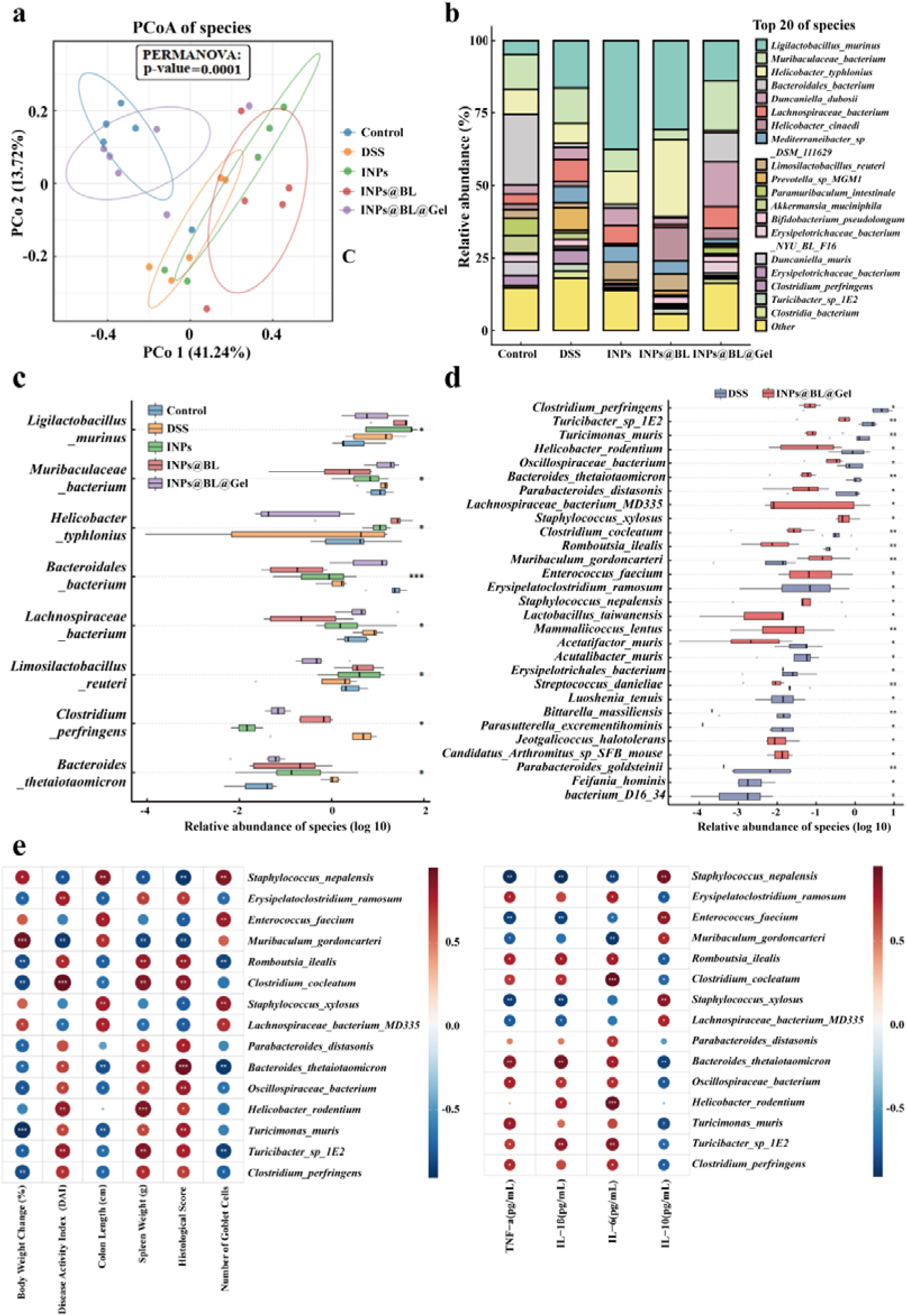
Regulation of INPs@BL@Gel on the homeostasis of intestinal microbiome. (a) Show the beta diversity of the intestinal microbiota of mice in different drug treatment groups through PCoA (PERMANOVA test). (b) The relative abundance of the Top 20 differential bacteria in the intestinal microbiota of each group of samples. (c) The relative abundance of bacteria with representative differences in each group of intestines (Kruskal-Wallis test). (d) Relative abundance of intestinal representative differential bacteria in DSS group and INPs@BL@Gel group (Kruskal-Wallis test). (e) Spearman correlation analysis between differential bacteria and inflammation-related penotypes and indicators. n = 5. *p<0.05, **p<0.01, ***p<0.001 and ****p<0.0001.

Although BL was not among the taxa exhibiting the largest intergroup differences, quantitative comparison revealed that BL abundance was significantly increased only in the INPs@BL@Gel group (Supplementary Figure S16). This finding indicates that the hydrogel-based formulation effectively enhances the survival and intestinal colonization of the administered probiotic strain. Pairwise comparison between the DSS and INPs@BL@Gel groups revealed selective suppression of inflammation-associated pathogenic taxa (Figure 9d). DSS-induced colitis resulted in pronounced enrichment of *Clostridium perfringens* and *Helicobacter rodentium*, both of which are known to exacerbate intestinal inflammation through toxin production and immune activation. In addition, a significant increase in *Bacteroides thetaiotaomicron* was observed in the DSS group. Although generally considered a commensal under physiological conditions, this species can adopt a mucin-degrading phenotype during inflammation, thereby compromising mucus barrier integrity. Notably, INPs@BL@Gel treatment reduced the abundance of these taxa to levels comparable to those in healthy controls, consistent with the restoration of epithelial and mucus barrier function limiting the ecological niche available to opportunistic pathobionts.

Concurrently, INPs@BL@Gel restored bacterial taxa that were depleted in DSS-induced colitis, including *Muribaculum gordoncarteri* and *Lachnospiraceae bacterium* MD335, both of which are recognized as SCFAs producers that support colonic energy metabolism and epithelial health. Spearman correlation analysis further linked these microbiota shifts to host phenotypic and inflammatory parameters (Figure 9e). DSS-enriched taxa, such as *C. perfringens* and *B. thetaiotaomicron*, showed positive correlations with disease severity indicators (DAI score, histological score, and spleen weight) and pro-inflammatory cytokines (IL-1β, IL-6, and TNF-α), while correlating negatively with body weight recovery and colon length. In contrast, taxa restored by INPs@BL@Gel, including *Muribaculum gordoncarteri* and *Lachnospiraceae bacterium MD335*, exhibited the opposite correlation pattern and were positively associated with the anti-inflammatory cytokine IL-10.

Overall, the metagenomic data demonstrate that INPs@BL@Gel induces targeted microbiome remodeling, characterized by suppression of mucin-associated pathobionts and enrichment of SCFAs-producing commensal bacteria. These microbiota changes are closely associated with improvements in intestinal inflammation and colitis-associated mental symptoms, supporting a microbiota-mediated mechanism underlying the therapeutic efficacy of INPs@BL@Gel.

### 2.10 Targeted Metabolite Analysis

To evaluate the functional consequences of microbiota remodeling, targeted metabolomic analyses were performed in both serum and brain tissues (For specific methods, please refer to the supporting materials). Hierarchical clustering of serum metabolites revealed pronounced metabolic dysregulation in DSS-treated mice (Figure 10a). Multiple metabolites, including HVA, butyrate, threonine, and histidine, were markedly reduced following DSS exposure, whereas these alterations were largely reversed upon INPs@BL@Gel treatment. Quantitative analyses further confirmed the restoration of microbially derived metabolites and essential nutrients in the circulation (Figure 10b). Among SCFAs, serum butyrate levels were significantly decreased in DSS mice (p < 0.05). Treatment with INPs@BL@Gel resulted in a clear upward trend in butyrate abundance; although statistical significance was not achieved due to inter-individual variability, mean values approached those of healthy controls. Previous studies have implicated butyrate in blood-brain barrier (BBB) repair and endothelial homeostasis, suggesting potential relevance to the observed neuroprotective effects.

**Figure 10.**
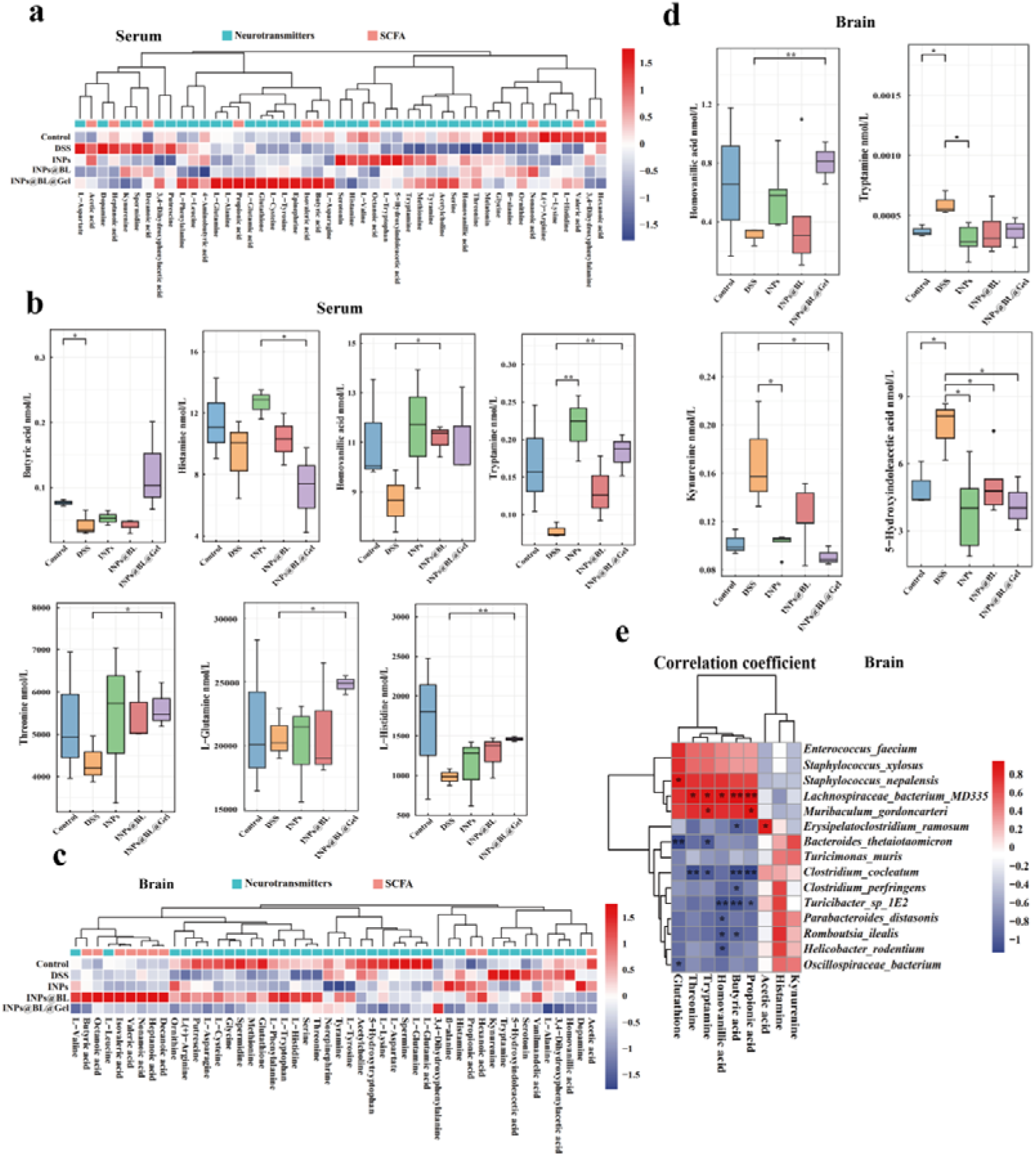
Targeted metabolomics analysis of serum and brain tissue after INPs@BL@Gel treatment. (a) Hierarchical clustering analysis of neurotransmitters and SCFAs in mouse serum following treatment with different drug groups, along with quantitative analysis of differential neurotransmitters and SCFAs (b). (c) Hierarchical clustering analysis of differential neurotransmitters and SCFAs in brain tissue across groups, and quantitative analysis of differential neurotransmitters and SCFAs (d). (e) Spearman correlation analysis between representative gut microbiota and differential neurotransmitter and SCFAs levels in brain tissue. Data are expressed as Mean ± SD, with n = 4.

We next focused on metabolites involved in neurotransmitter pathways. Serum HVA, the primary metabolite of dopamine, was significantly elevated in the INPs@BL@Gel group compared with the DSS group (p < 0.05), indicating recovery of peripheral dopamine metabolism. In addition, serum tryptamine—a microbiota-derived neuromodulator capable of crossing the BBB—was markedly depleted by DSS treatment but significantly restored following INPs@BL@Gel administration (p < 0.01). Consistent with these peripheral changes, brain metabolomic profiling revealed corresponding improvements in central neurochemical homeostasis. Heatmap analysis demonstrated recovery trends in several neuroactive precursors, as well as the antioxidant glutathione (Figure 10c). Quantitative analysis further showed that kynurenine levels were significantly elevated in the brains of DSS-treated mice (p < 0.05), consistent with activation of the indoleamine 2,3-dioxygenase (IDO) pathway (Figure 10d). INPs@BL@Gel treatment significantly reduced kynurenine levels (p < 0.05), thereby attenuating this neurotoxic metabolic route. Moreover, 5-hydroxyindoleacetic acid (5-HIAA), the major serotonin metabolite, was abnormally increased in DSS mice and was significantly normalized after treatment (p < 0.05), indicating correction of disrupted serotonin metabolism.

To link metabolic alterations with microbiota changes, Spearman correlation analysis was performed. *Lachnospiraceae bacterium* MD335 and *Muribaculum gordoncarteri*—both enriched following INPs@BL@Gel treatment—exhibited positive correlations with serum tryptamine and valeric acid levels (Figure 10e). The concurrent recovery of butyrate and tryptophan-derived metabolites provides a plausible mechanistic connection to BBB reinforcement. Butyrate, a potent histone deacetylase (HDAC) inhibitor, has been reported to enhance tight junction protein expression in brain endothelial cells through epigenetic regulation.^[42,43]^ Tryptamine, acting as a ligand for the aryl hydrocarbon receptor (AhR), contributes to vascular endothelial integrity.^[44,45]^ Together, these signaling pathways may reduce BBB permeability, limit the infiltration of peripheral pro-inflammatory cytokines, and suppress diversion of tryptophan toward the kynurenine pathway, thereby alleviating central neurochemical imbalance. Notably, HVA levels were increased in both serum and brain tissues following INPs@BL@Gel treatment. HVA has been associated with modulation of synaptic plasticity, particularly within prefrontal cortex and hippocampal circuits implicated in learning, emotion regulation, and cognitive recovery.^[21]^ Improved dopamine metabolism may therefore further support synaptic stability and remodeling by reducing oxidative stress and improving the neuroinflammatory microenvironment.

Collectively, the INPs@BL@Gel composite system exhibited sustained and comprehensive therapeutic efficacy against DSS-induced colitis and its associated neuropsychiatric comorbidities, supported by coordinated improvements in intestinal inflammation, behavioral phenotypes, barrier integrity, and gut microbiota homeostasis. In the DSS colitis model, INPs@BL@Gel restored colon length to 2.88-fold that of the DSS group mice, fully reversing the 33.04% shortening observed in DSS mice, and increased goblet cell abundance in colonic tissue by 20.3-fold, indicating effective repair of mucus barrier damage. In parallel, intestinal inflammation was markedly suppressed, with pro-inflammatory cytokines TNF-α, IL-1β, and IL-6 reduced by 33.9%, 61.31%, and 55.82%, respectively, while the anti-inflammatory cytokine IL-10 increased by 54.48%. Consistent with these molecular changes, histopathological injury scores decreased from 9 in DSS mice to 1 following treatment, approaching healthy control levels. Behavioral assessments demonstrated near-complete normalization of colitis-associated depressive- and anxiety-like behaviors. In the open-field test, treated mice traveled 43.83 m and spent 15.45% of the time in the center zone, effectively reversing the one-third reduction in locomotion and ∼50% decrease in center exploration induced by DSS. In the elevated plus maze, the proportion of time spent in open arms and the number of open-arm entries increased by 1.68-fold and 1.35-fold, respectively, compared with DSS mice. In the sucrose preference test, preference indices reached 0.74 at 3 h and 0.78 at 6 h, whereas DSS reduced these values by approximately one-third and two-fifths. Moreover, immobility times in the tail suspension and forced swim tests decreased by 37.9% and 62.96%, respectively, with no significant differences between treated and healthy control groups across all behavioral paradigms. Mechanistically, INPs@BL@Gel restored intestinal barrier function by upregulating ZO-1 and Occludin, eliminating excessive reactive oxygen species (91.0% clearance), and rescuing epithelial viability following H_2_O_2_ challenge from 74.5% to >90%. Concurrently, blood-brain barrier integrity was re-established, as evidenced by normalized Evans blue leakage, increased numbers of morphologically intact Nissl bodies in the hippocampal dentate gyrus and CA3 regions (1.84-fold and 4.28-fold vs. DSS), and enhanced synapsin-1 expression (7.64-fold and 8.59-fold, respectively). At the microbiota level, INPs@BL@Gel suppressed inflammation-associated taxa, including *Clostridium perfringens* and *Helicobacter rodentium*, while enriching SCFAs-producing commensals such as *Muribaculum gordoncarteri* and *Lachnospiraceae bacterium* MD335, with principal coordinates analysis demonstrating substantial convergence toward the healthy microbial profile, collectively indicating re-establishment of gut-brain axis homeostasis.

The overall therapeutic benefit of INPs@BL@Gel arises from a gut-brain dual-site intervention strategy delivered through two complementary routes: (i) targeted suppression of intestinal inflammation via a cascade of “site-specific release at inflamed regions-multicomponent synergy across pathological targets-microbiota remodeling”; and (ii) alleviation of neuropsychiatric symptoms through “complementary component functions-directed metabolite regulation-multitarget intervention.” INPs@BL@Gel is a composite hydrogel system centered on BL, rationally integrated with Bai, Inu, and Tyr to address both intestinal and neurological dysfunction. Following oral administration, the INPs@BL@Gel resists gastrointestinal degradation, enabling prolonged intestinal retention, stable bacterial survival, and effective colonization. Within inflamed intestinal regions, components of Bai, Inu, Tyr, and BL are locally released and act synergistically across multiple pathological targets to integrated therapy IBDs while reshaping the gut microbiota. In parallel, the system modulates intestinal metabolite profiles, increasing neuroactive metabolites such as HVA, tryptamine, and SCFAs (including butyrate and valerate) and leading to multi-target regulation of anxiety and depression, while reducing kynurenine and abnormally elevated 5-hydroxyindoleacetic acid (5-HIAA). Through gut-brain axis signaling, these coordinated metabolic shifts contribute to BBB repair and multi-target improvement of neuropsychiatric symptoms. Collectively, each component of INPs@BL@Gel performs multiple complementary functions, resulting in enhanced comprehensive therapeutic efficiency. This design concept and construction strategy provide experimental validation for engineered probiotic systems and gut-brain dual-site synergistic therapies.

## 3. Conclusion

In response to the clinical challenge posed by the frequent comorbidity of inflammatory bowel diseases (IBDs) with neuropsychiatric disorders such as depression and anxiety, this study proposes a “gut-brain dual-site, multi-target integrated synergistic and metabolite-regulated co-therapeutic strategy.” This concept is specifically designed to address the inherent limitations of conventional oral pharmacotherapy, which is constrained by the spatial separation of therapeutic targets in the intestine and brain, as well as the complex bidirectional regulation mediated by the gut-brain axis. This strategy was realized through the rational construction of the INPs@BL@Gel composite system. Bai (anti-inflammatory and neuroprotective), Tyr (precursor of HVA), Inu (prebiotic and promoter of SCFAs production), and BL were integrated into a coordinated polymeric network via Fe([) coordination to form INPs. Encapsulation within a dual-responsive hydrogel enabled inflammation-triggered release at colonic lesions, thereby establishing a stepwise therapeutic cascade consisting of multicomponent synergy, metabolic spectrum modulation and optimization, and ultimately gut-brain axis co-regulation. This system not only restored beneficial metabolites, such as normalizing HVA levels and elevating butyrate toward those of healthy controls, but also replenished essential amino acids (e.g., threonine and glutamine) required for intestinal mucosal repair. Collectively, this work introduces a new paradigm in probiotic engineering characterized by on-demand functional component integration and controllable metabolic outcome optimization.

By integrating baicalin and BL through multitarget coordination, the INPs@BL@Gel exploits the pH/MMP dual-responsive and acid-resistant properties of the SG-Gel to effectively protect probiotic viability. As a result, BL survival under gastric acid conditions increased by 56.87-fold, intestinal colonization was enhanced by 10.58-fold, colon length was restored by 2.88-fold, and goblet cell abundance increased by 20.3-fold, collectively demonstrating robust therapeutic efficacy in IBDs treatment. Importantly, by modulating key gut-derived metabolites—including HVA, SCFAs (such as butyrate and valerate), and amino acids (e.g., threonine and histidine)-the system also significantly alleviated colitis-associated depression. INPs@BL@Gel restored synapsin-1 (SYN1) expression in the hippocampal dentate gyrus to levels 8.59-fold higher than those observed in DSS-treated mice, reduced neurotoxic metabolites such as kynurenine, and reversed associated neuropathological alterations. Behavioral assays, including the open-field and tail suspension tests, together with neuromolecular analyses (SYN1 expression and Nissl body quantification), demonstrated that depression-like phenotypes in treated mice were largely indistinguishable from those of healthy controls.Furthermore, INPs@BL@Gel exhibited pronounced bidirectional barrier-protective effects, simultaneously preserving intestinal epithelial integrity and restoring blood-brain barrier (BBB) function. These findings provide compelling experimental evidence supporting the pathophysiological concept of “intestinal-derived mental disorders” and highlight the efficacy of INPs@BL@Gel in suppressing intestinal inflammation, ameliorating neuropsychiatric symptoms, restoring the barrier and re-establishing gut microecological homeostasis. Overall, this study presents a promising oral, long-acting co-therapeutic strategy for the treatment of colitis and its associated depression, laying a solid experimental foundation for future large-animal investigations and clinical translation.

## 4. Experimental Section

### Materials and Strains

Baicalin (Bai), ferric chloride (FeCl_3_), gelatin, calcium chloride (CaCl_2_), and *N,N*-dimethylformamide (DMF) were purchased from Aladdin (Shanghai, China). Tyrosine (Tyr), inulin (Inu), and polyethylene glycol (PEG 20000) were obtained from Macklin Biochemical Co., Ltd. (Shanghai, China). Dextran sulfate sodium (DSS; MW 36-50 kDa) was purchased from MP Biomedicals (California, USA). *Bifidobacterium longum* ATCC BAA-999 was obtained from the Ningbo Junyou Medical Technology Development Co., Ltd. Silkworm cocoons were provided by Jiangsu Fu’an Cocoon Silk Co., Ltd. (Yancheng, China). TPY liquid medium was purchased from Haibo Biotechnology Co., Ltd. (Qingdao, China). 2’,7’-Dichlorodihydrofluorescein diacetate (DCFH-DA) was obtained from Kaiji Biotechnology Co., Ltd. (Nanjing, China). Antibodies for immunofluorescence staining, includingPSD95 (RRID: AB_3102405), SYN1 (RRID: AB_10952778), occludin (RRID: AB_3070241), and ZO-1 (RRID: AB_3675805), were purchased from Hangzhou Hua’an Biotechnology Co., Ltd. (Hangzhou, China). All reagents were of analytical grade and used as received unless otherwise specified.

### Preparation of ICPs and INPs

DMF solutions of baicalin (5 mg mL^-1^) and FeCl_3_ (5 mg mL^-1^), as well as aqueous solutions of Tyr (0.45 mg mL^-1^) and Inu (5 mg mL^-1^), were prepared in advance. To synthesize the coordination polymer nanoparticles, 0.1 mL of the baicalin solution, 0.2 mL of the Tyr solution, and 0.2 mL of the FeCl_3_ solution were sequentially added to 10 mM Tris buffer containing Pluronic F127 (0.4 wt%). The mixture was stirred at room temperature for 1 h to allow coordination self-assembly, yielding Bai-Fe (III)-Tyr coordination polymer nanoparticles (ICPs). Subsequently, 0.2 mL of Inu solution was added to the ICPs dispersion and stirred continuously for an additional 4 h to form Inu covered Bai-Fe (III)-Tyr nanoparticles (INPs). The resulting nanoparticles were purified by dialysis, ultrafiltration, and centrifugation to remove unreacted components and solvents.

### Preparation of Silk Fibroin-Gelatin Composite Hydrogel (SG-Gel)

Silkworm cocoons were cut into small pieces and boiled in 0.5 wt% sodium carbonate solution to remove sericin, followed by thorough rinsing with deionized water and drying. The degummed silk fibroin (20 g) was dissolved in a ternary solvent system composed of CaCl_2_, ethanol, and deionized water (molar ratio 1:2:8) under heating and stirring until complete dissolution. The solution was centrifuged at 4500 rpm for 10 min, filtered, and dialyzed against deionized water to obtain a 2 wt% silk fibroin (SF) solution. The SF solution was then osmotically concentrated against 20 wt% PEG 20000 solution for 24 h to yield a 5 wt% SF solution.

Separately, a 5 wt% gelatin solution was prepared, and 1 wt% genipin solution was added as a crosslinker with thorough mixing. The concentrated SF solution was subsequently incorporated into the gelatin mixture, followed by incubation in a 37 °C water bath for 4 h to form the SG-Gel.

### Preparation of INPs@BL@Gel

BL was revived and cultured in TPY liquid medium to the stationary phase. One milliliter of bacterial suspension was collected, centrifuged, and washed twice with sterile phosphate-buffered saline (PBS). Subsequently, 200 μL of INPs solution (50 mg mL^-1^) was added to 1 mL of the BL suspension and gently mixed at room temperature for 1 h to allow nanoparticle association, yielding INPs@BL. The product was collected by centrifugation and washed twice with PBS to remove unbound nanoparticles. Finally, the INPs@BL suspension was resuspended in the SG-Gel precursor mixture and incubated at 37 [ for 4 h to obtain the composite formulation of INPs@BL@Gel.

### Characterization of INPs@BL@Gel

The hydrodynamic diameter and zeta potential of INPs were measured using a Malvern laser particle size analyzer (Zetasizer Nano ZS90, Malvern Instruments Ltd., UK). The morphology of the nanoparticles was examined by transmission electron microscopy (HRTEM, JEM-F200, JEOL, Japan). High-angle annular dark-field scanning transmission electron microscopy (HAADF-STEM, JEM-ARM200F, JEOL, Japan) was used to analyze the elemental distribution and composition of INPs@BL. The microstructures of SG-Gel and INPs@BL@Gel were observed by scanning electron microscopy (SEM, Sigma 360, Carl Zeiss, Germany). Ultraviolet-visible (UV-Vis) spectroscopy (T60, PG Instruments, China) and Fourier transform infrared spectroscopy (FTIR, TENSOR 27, Bruker, Germany) were employed to characterize the chemical composition of INPs and INPs@BL. High-performance liquid chromatography (HPLC, 1260 Infinity II, Agilent Technologies, USA) was used to quantify the loading content and relative ratios of active components in the nanoparticles. Chromatographic separation was performed using acetonitrile as mobile phase A and 0.5% phosphoric acid aqueous solution as mobile phase B under gradient elution conditions. X-ray photoelectron spectroscopy (XPS, ESCALAB Xi[, Thermo Fisher Scientific, USA) was used to analyze the surface chemical composition of INPs. X-ray diffraction (XRD, D8 ADVANCE, Bruker, Germany) was conducted to evaluate their crystalline structure. Fluorescence images were acquired using a super-resolution laser scanning confocal microscope (TCS SP8 STED 3X, Leica Microsystems, Germany) and a panoramic tissue and cell quantitative analysis system (Axioscan-7, Carl Zeiss Microscopy, Germany).

### Cell Lines and Culture Conditions

Mouse hippocampal neuronal cells (HT 22, RRID: CVCL_0321) and human normal colon epithelial cells (NCM 460, RRID: CVCL_0460) were obtained from ProNova Biotechnology Co., Ltd. (Wuhan, China). Both HT22 and NCM 460 cells were cultured in Dulbecco’s Modified Eagle Medium (DMEM) supplemented with 10% fetal bovine serum. All cells were maintained at 37 °C in a humidified incubator with 5% CO_2_.

### External Environmental Resistance and Responsive Release Behavior of INPs@BL@Gel

Simulated gastric fluid (SGF) and simulated intestinal fluid (SIF) were prepared according to previously reported protocols.^[30]^ Equal amounts (1 × 10^8^ CFUs) of BL, INPs@BL, and INPs@BL@Gel were incubated in 1 mL of SGF or SIF, respectively. At predetermined time points, aliquots of bacterial suspensions were collected, serially diluted, and plated on TPY agar plates. After incubation at 37 °C for 48 h under anaerobic conditions, colony-forming units were counted to assess bacterial viability.

To evaluate enzyme-responsive degradation and bacterial release, INPs@BL@Gel was incubated in 2 mL of SIF in the presence or absence of matrix metalloproteinases (MMPs). At designated time points, changes in hydrogel morphology were visually recorded, and the remaining hydrogel mass was weighed. Concurrently, released bacterial suspensions were collected, serially diluted, and plated onto TPY agar plates. After 48 h of anaerobic incubation, bacterial colonies were enumerated.

### Mucosal Adherence Capacity of INPs@BL@Gel

INPs were fluorescently labeled, and equal amounts of fluorescently labeled INPs, INPs@BL, and INPs@BL@Gel were co-incubated with freshly isolated mouse colon tissue at 37 °C for 1 h. After incubation, the tissues were washed twice with PBS to remove non-adherent materials and imaged by fluorescence microscopy.

In vivo mucosal adhesion and intestinal retention were further evaluated using a small-animal fluorescence imaging system (AMI HTX, Spectral Instruments Imaging, USA). DSS-induced colitis mice (female C57BL/6, 6-8 weeks old) were randomly assigned to three groups (n = 3) and orally administered equal volumes of INPs, INPs@BL, or INPs@BL@Gel. Fluorescence signals were recorded at predetermined time points in live animals and in excised colon tissues.

All animal experiments were conducted using C57BL/6 mice purchased from the Experimental Animal Center of Xi’an Jiaotong University School of Medicine and were approved by the Ethics Committee of Xi’an Jiaotong University (Approval No. 201929).In addition, all procedures were performed in accordance with *the Guide for the Care and Use of Laboratory Animals* and relevant regulations/guidelines. To minimize animal pain and distress, all procedures that could potentially cause pain or stress were conducted under appropriate anesthesia; throughout the study, trained personnel provided food daily and maintained animals under appropriate temperature conditions.

### ROS-Scavenging Ability of INPs@BL@Gel

(1) ABTS Radical Scavenging Assay: The ABTS radical cation (ABTS•[) was generated by mixing ABTS solution (7.4 mM) with potassium persulfate (K_2_S_2_O_8_, 2.6 mM) and incubating the mixture in the dark for 2 h at room temperature. The activated ABTS•[ solution was then mixed with INPs@BL@Gel at different concentrations. After incubation, the absorbance at 734 nm was measured using a UV-Vis spectrophotometer, with anhydrous ethanol serving as the blank control.^[46]^ The ABTS scavenging activity was calculated as: ABTS scavenging capacity = (A_0_ - A)/A_0_ × 100%. where A_0_ is the absorbance of the control and A is the absorbance of the sample. (2) DPPH Radical Scavenging Assay: A DPPH solution (0.05 mg mL^-1^) prepared in anhydrous ethanol was mixed with INPs@BL@Gel at varying concentrations. After incubation, the absorbance at 519 nm was recorded, with anhydrous ethanol as the control.^[46]^ The DPPH scavenging capacity was calculated as: DPPH scavenging capacity = (A_0_ - A)/A_0_ × 100%. (3) Hydroxyl Radical (•OH) Scavenging Assay: The hydroxyl radical scavenging ability was evaluated using a Fenton reaction system. Briefly, INPs@BL@Gel at different concentrations was added to a reaction mixture containing FeSO_4_ (6 mM), salicylic acid (6 mM), and H_2_O_2_ (8.8 mM). The mixture was incubated at 37°C for 30 min, after which the absorbance at 510 nm was measured.^[47]^ The •OH scavenging capacity was calculated as: •OH scavenging capacity = (A_1_ - A_2_))/A_0_ × 100%. (4) Superoxide Anion (O[•[) Scavenging Assay: INPs@BL@Gel solutions (0.8 mL) at different concentrations were mixed with 0.8 mL of 50 mM Tris-HCl buffer and incubated at 25 °C for 20 min. Subsequently, 0.4 mL of xanthine solution (1.5 mM) was added to initiate superoxide generation. The absorbance at 320 nm was recorded spectrophotometrically.^[47]^ O_2_•[ scavenging capacity = (A_1_ - A_2_))/A_0_ × 100%. (5) Intracellular ROS-Scavenging Capacity: Human normal colonic epithelial cells (NCM 460) were seeded into 96-well plates and cultured for 24 h. Cells in the control group received fresh medium only, whereas cells in the oxidative stress groups were treated with either 100 μM H_2_O_2_ or 50 ng mL[¹ lipopolysaccharide (LPS). INPs, INPs@BL, or INPs@BL@Gel were then added to the respective wells and incubated at 37 °C for 24 h. After treatment, the culture medium was removed and replaced with serum-free medium containing 10 μM 2’,7’-dichlorofluorescein diacetate (DCFH-DA). Cells were incubated at 37 °C in the dark for 30 min, followed by three washes with serum-free medium to remove excess probe. Intracellular ROS levels were subsequently evaluated by laser confocal microscopy and quantified by flow cytometry.

### Establishment and Treatment of DSS-Induced Colitis-Associated Mental Disorder Model

Female C57BL/6 mice (6-7 weeks old) were acclimated for 7 days prior to experimentation. Colitis-associated depression was induced by administering drinking water containing 3% (w/v) dextran sulfate sodium (DSS) for 7 consecutive days. Healthy control mice received regular drinking water.

After DSS induction, mice were randomly assigned to four treatment groups (n = 5 per group) and orally administered PBS, INPs, INPs@BL, or INPs@BL@Gel for 5 consecutive days. Following treatment, mice were euthanized, and fecal samples and tissues were collected for subsequent analyses.

### Evaluation of BBB Integrity and Permeability

Transmission Electron Microscopy (TEM): Mice were anesthetized with 10% sodium pentobarbital and transcardially perfused with sterile PBS to remove circulating blood. This was followed by perfusion with a fixative solution containing 0.25% glutaraldehyde and 4% paraformaldehyde for 5-10 min. Brain tissues from the prefrontal cortex were harvested and further fixed in 2.5% glutaraldehyde and 2% paraformaldehyde at 4 °C for 2 h.

The tissues were rinsed with phosphate-buffered saline, post-fixed in 1% osmium tetroxide, dehydrated through a graded ethanol series, embedded, sectioned into ultrathin slices, and stained with uranyl acetate and lead citrate. BBB ultrastructure-including endothelial cells, basement membranes, and astrocytic end-feet-was examined using TEM.

### Evans Blue (EB) Extravasation Assay

BBB permeability was assessed using the Evans Blue dye method. Briefly, mice received a tail-vein injection of 80 μL of 2% (w/v) EB solution. After 1 h, mice were perfused transcardially with PBS to remove intravascular dye. Brain tissues were harvested, photographed, weighed, and homogenized in DMSO. Samples were incubated at 60 °C for 24 h to extract EB, followed by centrifugation. The absorbance of the supernatant was measured at 620 nm to quantify EB extravasation.

### Behavioral Assessments

All behavioral tests were performed during the dark phase under low-light conditions (<20 lx). Mouse movements were tracked and analyzed using ANY-maze 7.3 software. To minimize stress-induced variability, mice were gently handled daily for three days prior to behavioral testing.

### Open Field Test (OFT)

The open field test was conducted to assess locomotor activity and exploratory behavior.^[48]^ Mice were placed individually in a 50 × 50 × 40 cm open-field arena and allowed to explore freely for 10 min. Behavioral parameters—including total distance traveled, movement trajectories, and time spent in the central zone—were recorded using infrared night-vision cameras and analyzed with ANY-maze 7.3 software. The apparatus was cleaned with 75% ethanol between trials.

### Elevated Plus Maze (EPM)

The elevated plus maze test was used to evaluate anxiety-like behavior.^[48]^ The apparatus consisted of two open arms and two closed arms elevated 60 cm above the floor. Mice were placed in the central area and allowed to explore freely for 10 min. Time spent in open arms, number of open-arm entries, and locomotor activity were recorded and analyzed using ANY-maze 7.3. The maze was cleaned with 75% ethanol after each test.

### Sucrose Preference Test (SPT)

The sucrose preference test was used to assess anhedonia-like behavior.^[48]^ Mice were housed individually and provided with two identical bottles containing either 1% sucrose solution or plain drinking water. Bottle positions were alternated every 12 h to avoid location bias. After a 24 h food and water deprivation period, sucrose and water consumption were measured at 3 h and 6 h intervals. Sucrose preference was calculated as the ratio of sucrose intake to total fluid intake.

### Tail Suspension Test (TST)

The Tail Suspension Test (TST) is widely used to evaluate depressive-like behaviors, including behavioral despair and anhedonia, in experimental animals.^[48]^ Briefly, a 60 cm-high stand was placed in a quiet testing environment. The distal end of the mouse tail (approximately 1 cm from the tip) was securely attached with adhesive tape, and the opposite end of the tape was fixed to the stand, allowing the mouse to be suspended in an inverted position with its head maintained at a safe distance above the floor. Each mouse was visually isolated from others using opaque partitions. The total test duration was 6 min, with the first minute defined as an adaptation period during which mice exhibited vigorous struggling to adjust to the abnormal posture. Immobility time was recorded throughout the test using infrared night-vision cameras and analyzed with the animal behavior analysis system ANY-maze 7.3. Upon completion of the test, mice were gently removed from the apparatus, the adhesive tape was detached, and animals were returned to their home cages.

### Forced Swimming Test (FST)

The Forced Swimming Test (FST) is a classical behavioral despair paradigm that exploits rodents’ innate aversion to water.^[48]^ When placed in an inescapable aquatic environment, mice initially engage in active escape-directed behaviors, followed by the gradual emergence of immobility, indicative of behavioral despair. For this assay, a 2 L glass beaker was filled with water maintained at approximately 25°C. The water depth was adjusted according to body weight to ensure that the mouse’s tail did not touch the bottom of the beaker. Each mouse was gently introduced into the water along the beaker wall and allowed to swim for a total of 8 min. The first 2 min were considered an adaptation phase, and behavioral parameters—including swimming time, immobility time, and climbing time—were recorded during the remaining 6 min. After testing, mice were promptly removed from the water, dried with a warm air stream, and returned to their cages.

### Metagenomic Sequencing and Analysis

Stool samples were submitted to Suzhou Shanjun Biomedical Technology Co., Ltd. (Suzhou, China) for metagenomic analysis. DNA extraction was performed using the Magbeads Fast DNA kit, followed by library construction with the KAPA HyperPlus kit and shotgun sequencing on the MGISEQ-2000 platform. Raw sequencing reads were quality-filtered using fastp (v0.23.0), and host-derived reads were removed by alignment against the GRCm39 reference genome using Bowtie2 (v2.3.5.1). Taxonomic profiling was conducted with MetaPhlAn4, while functional pathway annotation was performed using HUMAnN3 based on KEGG, MetaCyc, and gene family models (GMMs). Alpha diversity indices (Shannon and Simpson) and beta diversity metrics (Bray-Curtis distance with principal coordinates analysis) were compared using the Kruskal-Wallis test and PERMANOVA, respectively. Differential taxa and metabolic pathways were identified using LEfSe analysis with an LDA score threshold > 2.0. Statistical analyses were conducted in R software (v4.3.1) using Wilcoxon rank-sum and Fisher’s exact tests, with false discovery rate (FDR)-corrected p values < 0.05 considered significant.

### Statistical Analysis

Unless otherwise specified, data are presented as mean ± standard deviation (SD) from three independent experiments. Statistical differences between two groups were analyzed using Student’s t-test, while comparisons among multiple groups were performed using one-way analysis of variance (ANOVA) followed by a least significant difference (LSD) post hoc test where appropriate. A P value < 0.05 was considered statistically significant.

## Acknowledgements

Zhang S, Zhang Y, and He J contributed equally to this study. This work was funded by the National Natural Science Foundation of China (No. 82271572&NO. 82230044), National Key R&D Program of China (Grant No. 2025YFC2511200). China Postdoctoral Science Foundation (2025M782947). This investigation was supported by the Institute of Shaanxi Provincial Key Laboratory of Biological Psychiatry, the First Affiliated Hospital of Xi’an Jiaotong University. And we thank the Instrument Analysis Center of Xi’an Jiaotong University for their assistance with the measurements.

## Data Availability Statement

The data that support the findings of this study are available from the corresponding author upon reasonable request.

