## Supplementary material for "Multi-component functionalized *Bifidobacterium longum* hydrogel for multi-target integrated therapy of colitis-associated anxiety and depression": -

Zhang S^1^, Ma Q^1^, Ma S^1^, Li C^1^, Ma X^1,2^, Zhu F^1,2^

Center for Brain Science & Department of Psychiatry, The First Affiliated Hospital of Xi’an Jiaotong University, Xi’an 710061, China

Zhang Y^2^, Wang Y^2^, Jin S^2^, Ma X^1,2^, Zhu F^1,2^

Center for Translational Medicine & Department of Psychiatry, The First Affiliated Hospital of Xi’an Jiaotong University, Xi’an 710061, China

He J^3^, Li S^3^, Li Q^3^, Xie X^3^, Zhang H^3^, Deng J^3^, Wu D^3^

The Key Laboratory of Biomedical Information Engineering of Ministry of Education, School of Life Science and Technology, Xi’an Jiaotong University, Xi’an, 710049, China

Zhang Y^4^

Department of Breast Disease,The Affiliated Cancer Hospital of Zhengzhou University and Henan Cancer Hospital, Zhengzhou, 450008, China

Song X^5^

Biological Psychiatry International Joint Laboratory of Henan, Zhengzhou University, Zhengzhou, 450052, China

^#^ These authors contributed equally to this work

**Supplementary Material 1：Experimental method**

**Chromatographic and Mass Spectrometric Conditions:** The mobile phase consisted of 0.01% formic acid in water (phase A) and 1 mM ammonium formate in 95% methanol (phase B). The column temperature was maintained at 45 ℃, the autosampler temperature at 6 ℃, and the injection volume was set to 1 μL. Mass spectrometric detection was performed using a 6500 QTRAP+ triple quadrupole mass spectrometer equipped with an electrospray ionization (ESI) source operating in multiple reaction monitoring (MRM) mode. Data acquisition and processing were carried out using Analyst software (v1.7.3, SCIEX) and Biotree Biobud (v2.0.3).

**Targeted Metabolomics Analysis:** Targeted quantitative analyses of short-chain fatty acids (SCFAs) and neurotransmitters were performed by BioQuad Co., Ltd. Fecal samples were transferred into 2 mL EP tubes, vortexed, homogenized using a ball mill for 4 min, and subjected to ultrasonication for 5 min in an ice-water bath, repeated three times. Samples were centrifuged at 5000 rpm and 4 ℃ to collect the supernatant. Subsequently, 0.1 mL of 50% H_2_SO_4_ and 0.8 mL of extraction solution containing 25 mg/L methyl tert-butyl ether as an internal standard were added. The mixture was vortexed, shaken, sonicated for 10 min (ice-water bath), and centrifuged at 10,000 rpm for 15 min at 4 ℃. Samples were then stored at -20 ℃ for 30 min, and the supernatant was transferred to 2 mL glass vials for GC-MS analysis. Serum samples were processed similarly by adding 0.05 mL of 50% H_2_SO_4_ and 0.2 mL of extraction solution containing the internal standard, followed by vortexing, ultrasonication, centrifugation, and cold storage prior to GC-MS analysis. For neurotransmitter quantification, serum and brain tissue samples were extracted using pre-chilled acetonitrile containing 0.1% formic acid. Serum samples (20 μL) were mixed with 80 μL of extraction solvent, vortexed, and sonicated. Brain tissue samples were weighed, combined with copper beads, 80 μL of extraction solvent, and 20 μL of water, followed by repeated grinding and ultrasonication in an ice-water bath. All samples were incubated at −20 °C overnight for protein precipitation, centrifuged at 12,000 rpm for 15 min at 4 °C, and derivatized with Na_2_CO_3_ and L-BzCl reagent. After the addition of the internal standard (H-std-a-Bz-DF) and centrifugation, supernatants were collected for analysis using an Agilent 1290 Infinity UHPLC system equipped with an ACQUITY UPLC BEH C18 column (1.7 μm, 2.1 mm × 150 mm).

**Supplementary Material 2：**

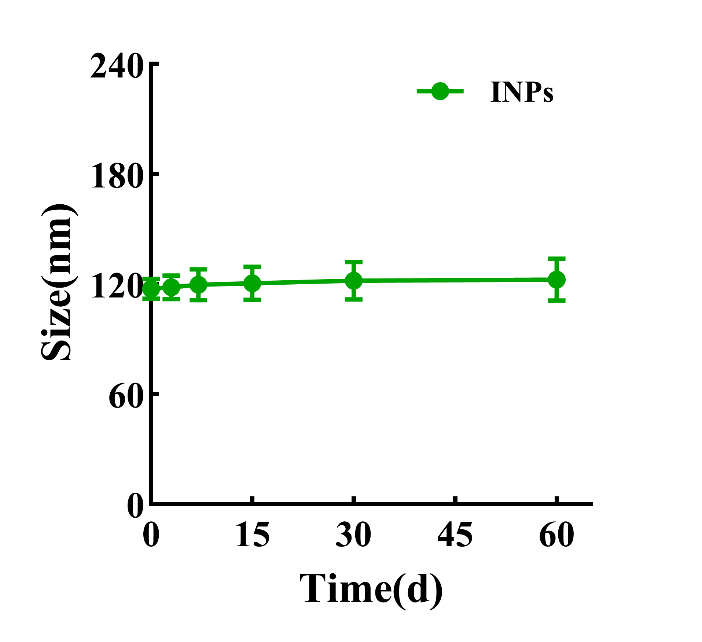

**Figure S1.** Particle size changes in INPs over 60 days in a medium containing 10% FBS.

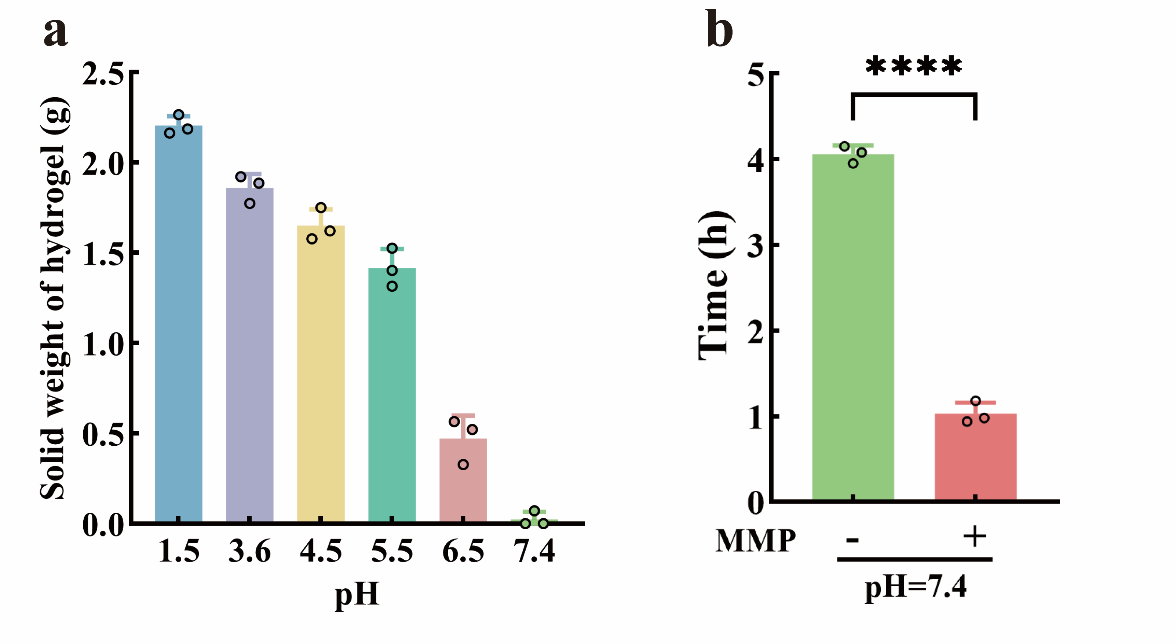

**Figure S2.** The pH- and MMP-Responsive Release from SG-Gel. (a) Change in Residual Solid Weight of SG-Gel After 4h Under Different pH Conditions. (b) Time required for complete dissolution of SG-Gel at pH = 7.4 in the presence/absence of MMP. Data are expressed as Mean ± SD, with n = 3. Statistical analysis was evaluated with two-tailed Student’s t tests (*p < 0.05, **p < 0.01, ***p < 0.001, and ****p < 0.0001).

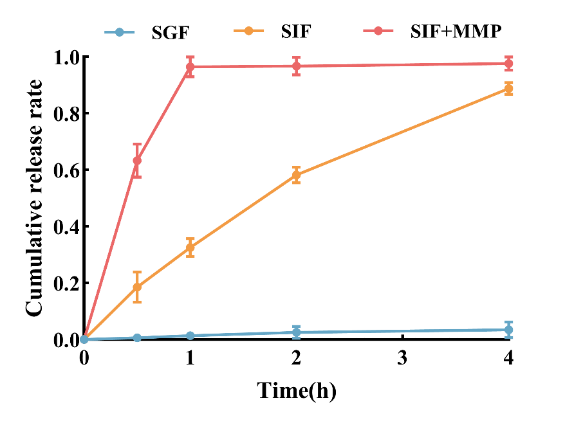

**Figure S3.** Cumulative release profiles of samples over 4 hours in simulated gastric fluid (SGF), simulated intestinal fluid (SIF), and MMP-containing simulated intestinal fluid (SIF + MMP). Data are expressed as Mean ± SD, with n = 3.

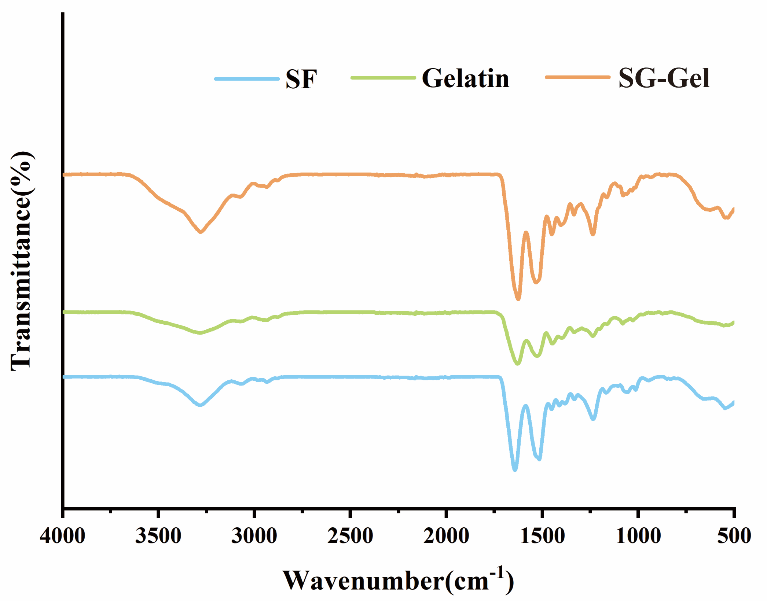

**Figure S4.** FTIR of SF, Gelatin and SG-Gel.

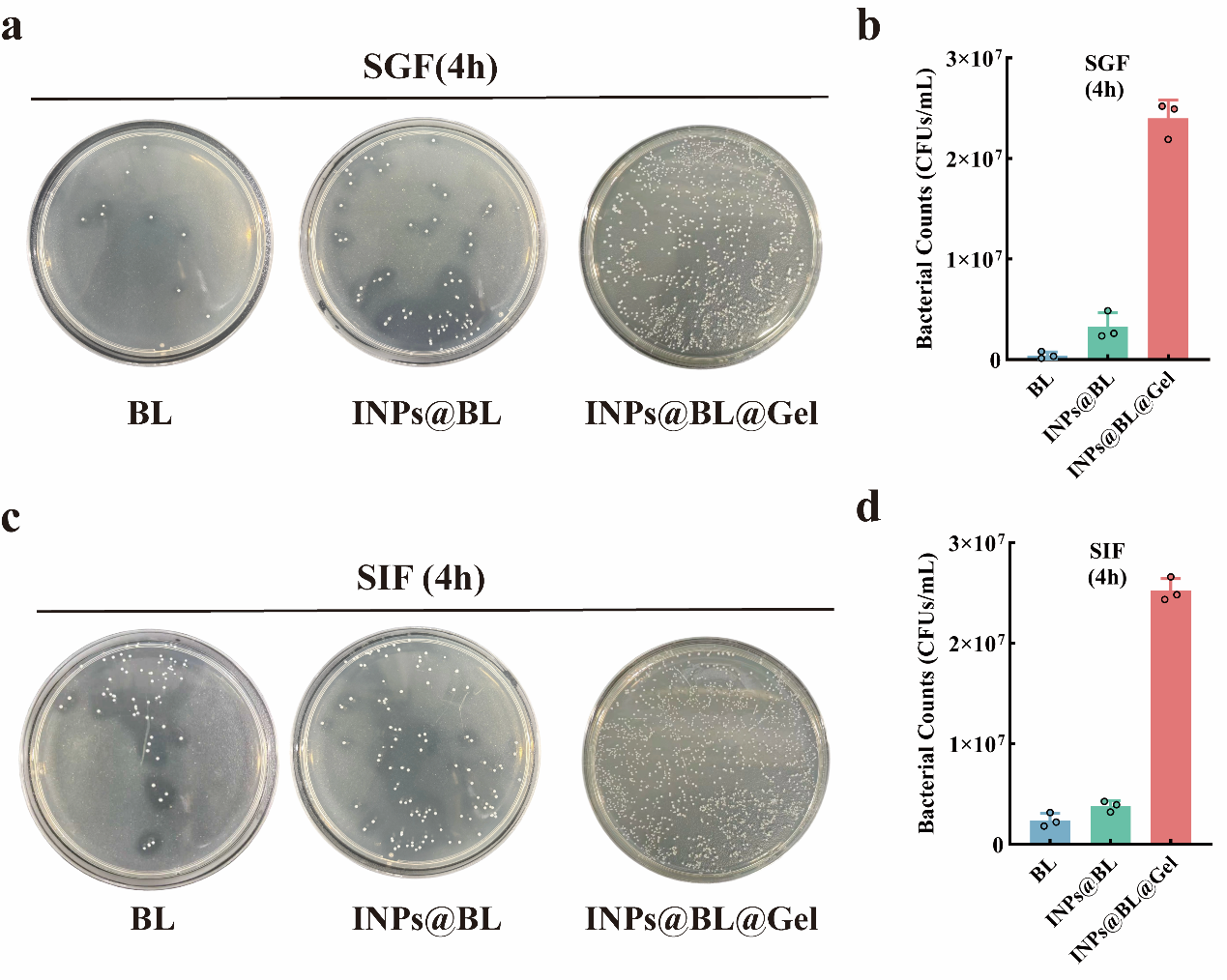

**Figure S5.** Bacterial counts of INPs@BL@Gel after digestion in simulated gastric fluid (SGF) and simulated intestinal fluid (SIF). (a, b) Photographs and corresponding counts of bacterial colonies on TPY AGAR plates in simulated gastric fluid (SGF); (c, d) Photographs and corresponding counts of bacterial colonies on TPY AGAR plates in simulated intestinal fluid (SIF). Data are expressed as Mean ± SD, with n = 3.

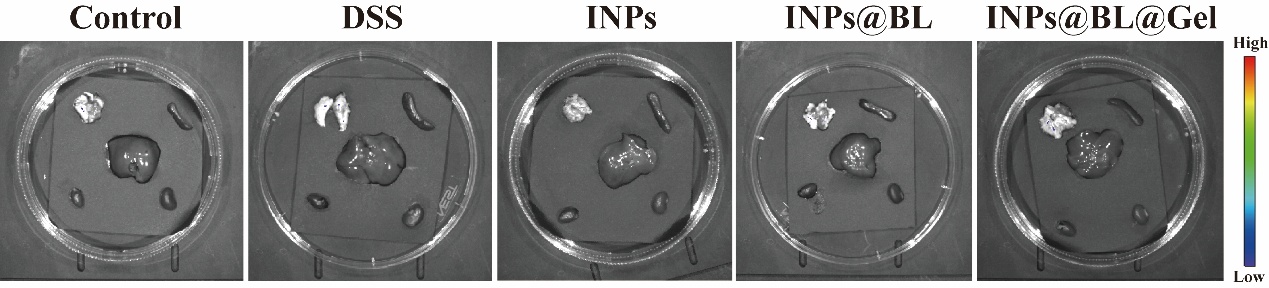

**Figure S6.** Representative fluorescence images of the heart, liver, spleen, lungs, and kidneys in mice from different drug groups labeled with IR780 after oral administration.

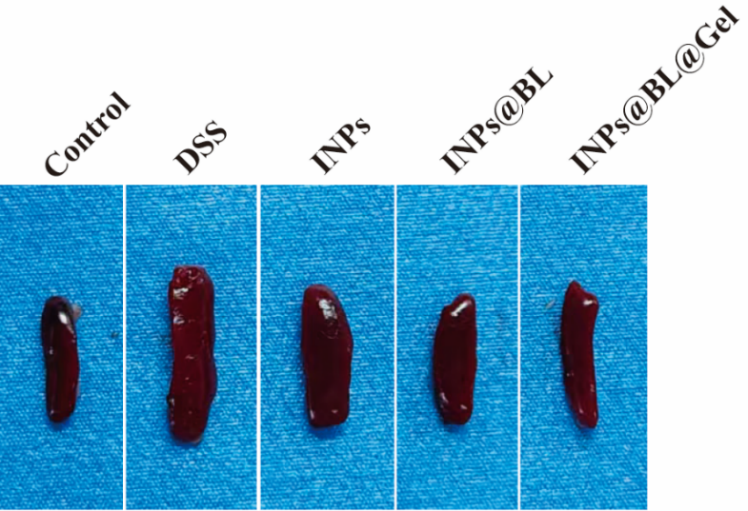

**Figure S7.** Representative pictures of the spleens of mice in different groups. The INPs@BL@Gel group showed no significant splenomegaly, comparable to the control group.

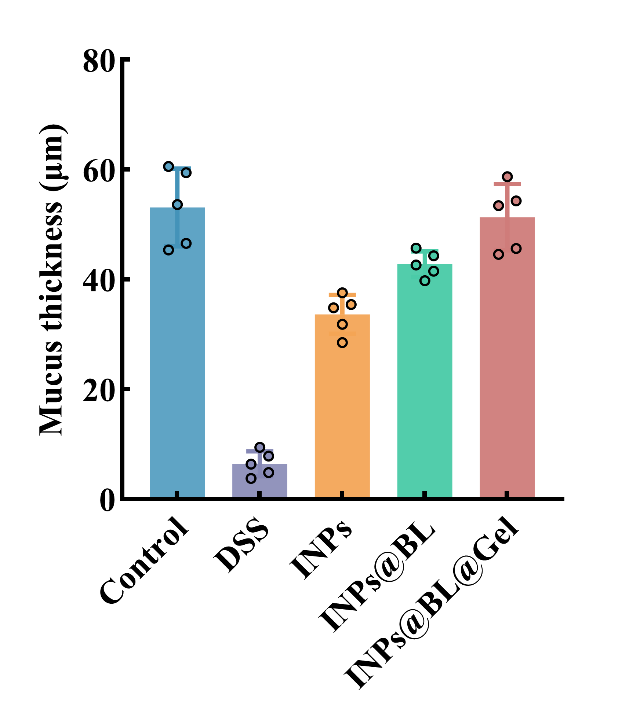

**Figure S8.** Assessment of mucus thickness in the colons of mice in each group by AB-PAS staining. Data are expressed as Mean ± SD, with n=5.

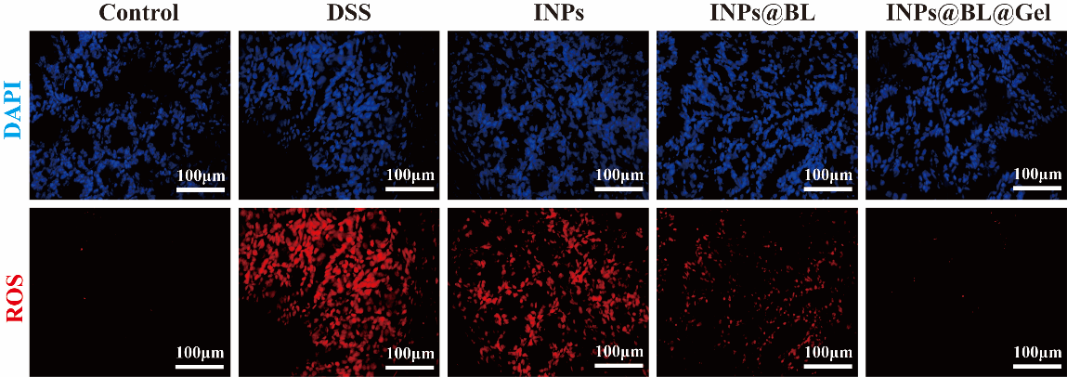

**Figure S9.** Representative ROS immunofluorescence staining images of mouse colons from each group. Cell nuclei were stained with DAPI (blue).

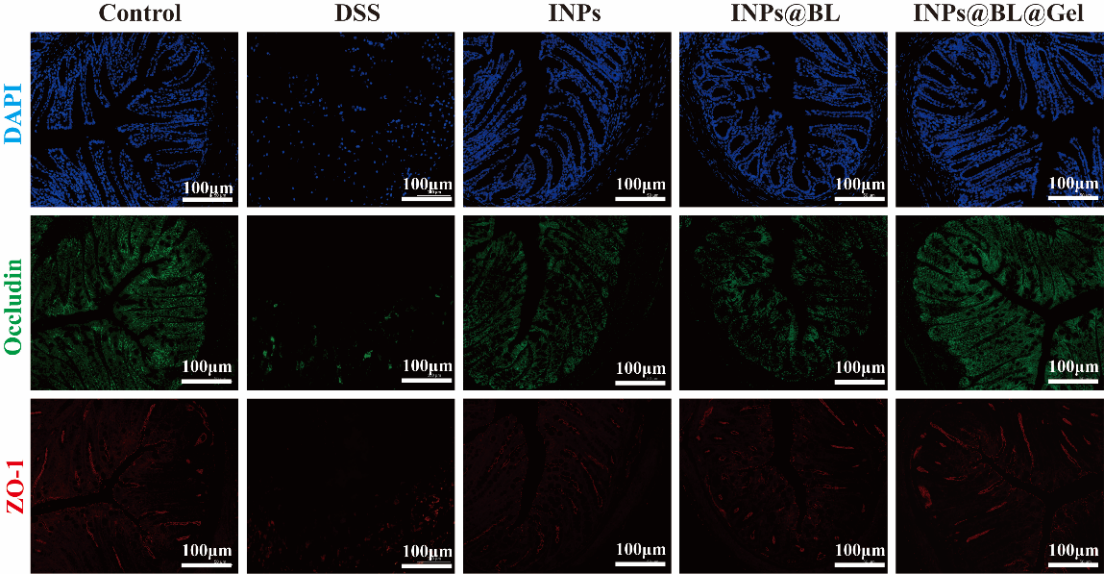

**Figure S10.** Representative immunofluorescence staining images of tight junction proteins ZO-1 (red) and occludin (green) in the colons of each group of mice. Cell nuclei were stained with DAPI (blue).

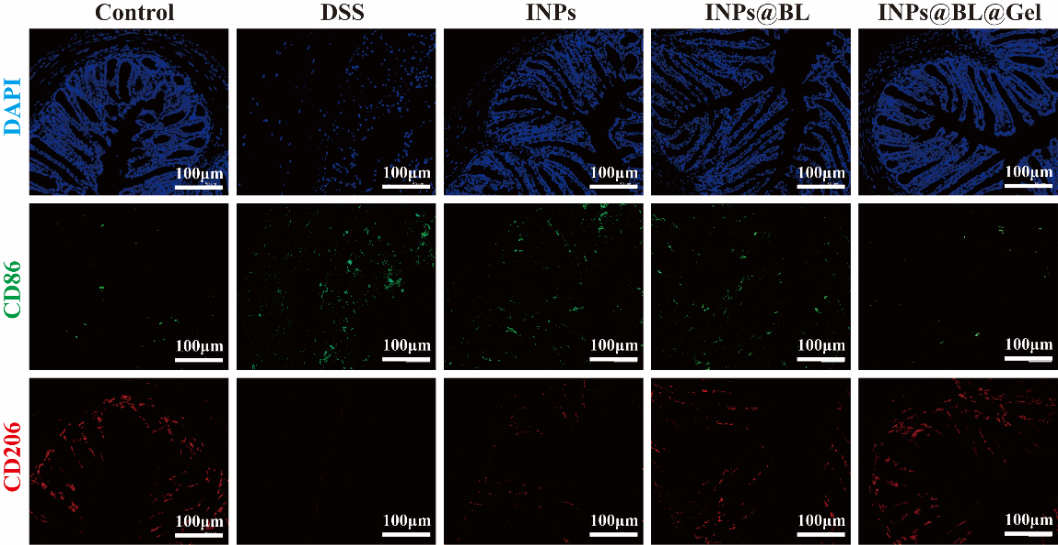

**Figure S11.** Representative immunofluorescence staining images of CD86 (green), a marker for M1 macrophages, and CD206 (red), a marker for M2 macrophages, in the colons of mice from each group. Cell nuclei were stained with DAPI (blue).

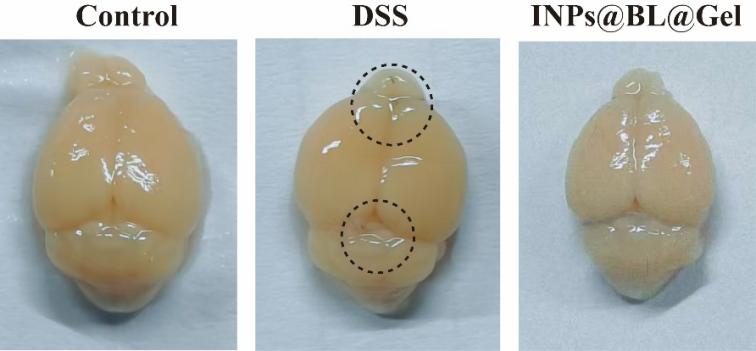

**Figure S12.** Representative images of blood-brain barrier (BBB) permeability in Control, DSS, and INPs@BL@Gel groups assessed via intravenous injection of Evans Blue (EB).

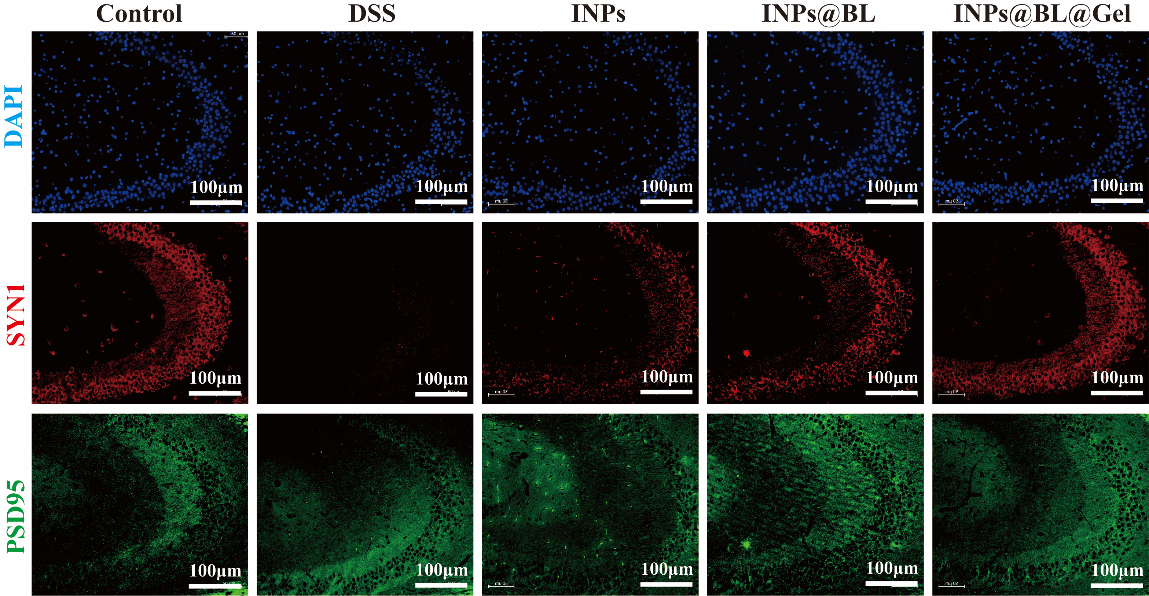

**Figure S13.** Representative immunofluorescence staining images of synapse-associated proteins SYN1 (red) and PSD95 (green) in the hippocampal CA3 region of mice from each group. Cell nuclei were stained with DAPI (blue).

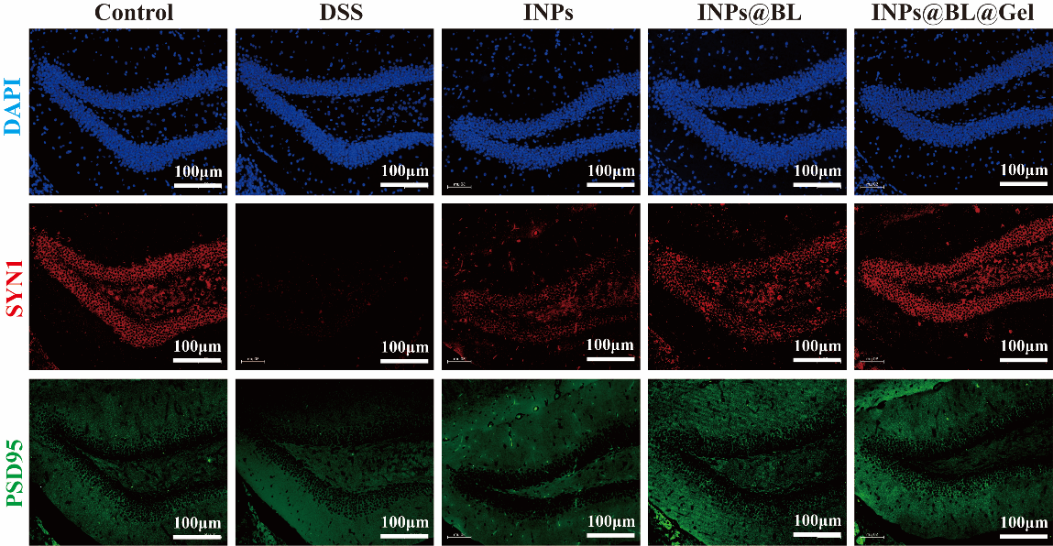

**Figure S14.** Representative immunofluorescence staining images of synapse-associated proteins SYN1 (red) and PSD95 (green) in the hippocampal DG region of mice from each group. Cell nuclei were stained with DAPI (blue).

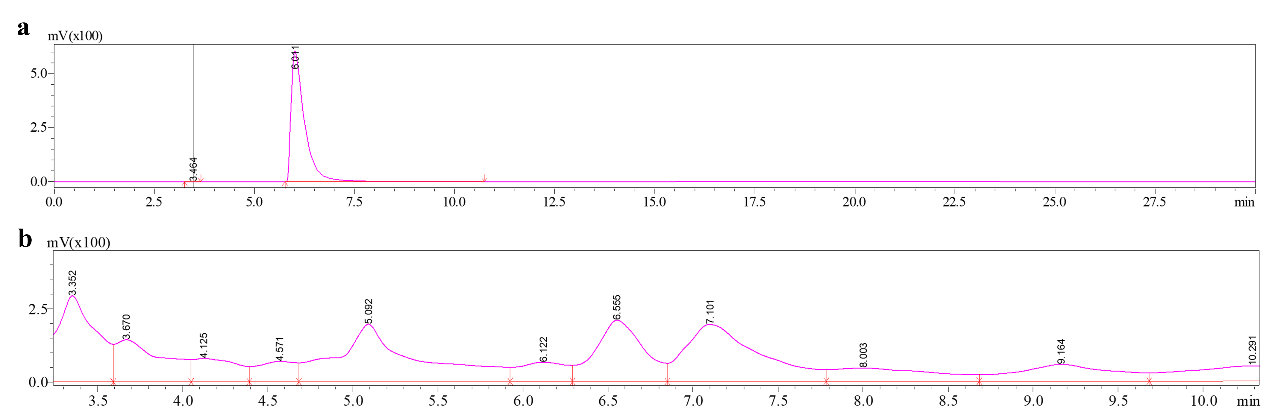

**Figure S15.** HPLC Chromatographic Analysis of HVA Standards (a) and INPs@BL@Gel Samples (b).

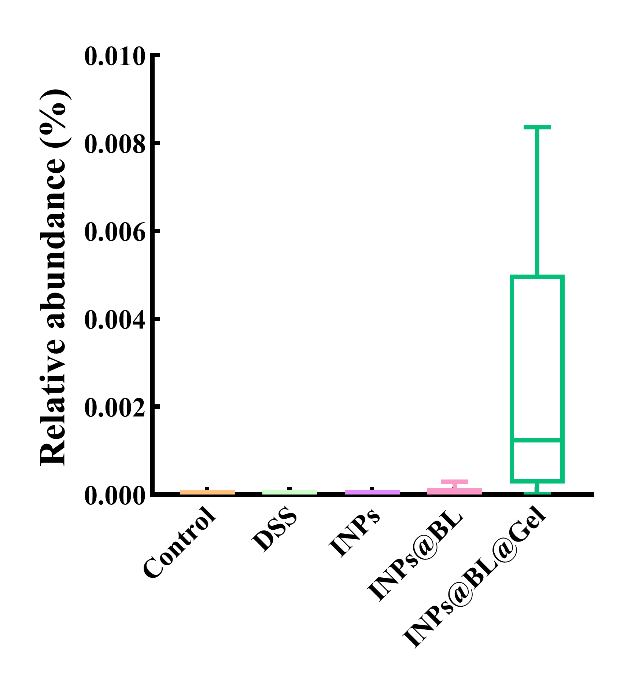

**Figure S16.** Relative abundance of BL in the feces of each group of mice. Data are expressed as Mean ± SD, with n = 5.

**Supplementary Material 3：**

**Table S1.** The electron binding energy of coordination atoms in Fe(III)-Bai-ICP NPs and its"preparationraw"materials.

| Sample | Binding energy[eV] | | | | | | |
| --- | --- | --- | --- | --- | --- | --- | --- |
|  | O1s | | | Fe 2p 3/2 | | Fe 2p 1/2 | |
| FeCl_3_ | - | | | 711.80 eV | | 725.16 eV | |
| Bai | 529.30 eV | 53 1.20 eV | - | | - | - | - |
| Fe(III)-Bai ICPs | 531.30 eV | 532.20 eV | 709.10 eV | | 711.58 ev | 721.90 eV | 724.20 eV |

**Table S2.** Scoring system of DAI

| Score | Body weight loss | Stool consistency | Blood |
| --- | --- | --- | --- |
| 0 | ≤1% | Normal | Negative hemoccult |
| 1 | 1-5% | Soft but formed |  |
| 2 | 5-10% | Soft | Positive hemoccult |
| 3 | 10-15% | Very soft |  |
| 4 | >15% | Watery diarrhea | Blood traces in stool visible |

DAI score = (Body weight loss score + Stool consistency score + Blood score)/3

**Table S3.** Histopathological Assessment of Colitis

| Feature | Score | Description |
| --- | --- | --- |
| Mucosal epithelium | 0 | No mucosa inflammation/prolonged epithelialcells |
|  | 1 | Destruction of barrier/<10% loss of epithelialsurface ulcer |
|  | 2 | 10-30% loss of epithelial surface ulcer |
|  | 3 | 30-60% loss of epithelial surface ulcer |
|  | 4 | >60% loss of epithelial surface ulcer |
| Crypt | 0 | No mucosa inflammation/intact crypts |
|  | 1 | Destruction of barrier/<10% loss of cryptsulcer |
|  | 2 | 10-20% loss of crypts ulcer |
|  | 3 | >20% loss of epithelial surface and crypts |
| Cell infiltration and edema | 0 | None |
|  | 1 | Mild Infiltration |
|  | 2 | Moderate Infiltration |
|  | 3 | Severe Infiltration |
| Goblet cells depletion | 0 | Absent |
|  | 1 | Present |

**Table S4.** Nerve injury scores of Brain.

| Degree of injury | Description | Score |
| --- | --- | --- |
| Normal | Cells and extracellular matrix were uninjured | 0 |
| Mildly injured | Slightly swollen cells with decreased protrusions spread in the homogenousextracellular matrix. Pyknotic cells, apoptosis or necrosis was rarely observed | 1 |
| Moderately injured | Oval, lightly stained swollen cells. Increased red neurons, pyknotic cells,apoptosis and necrosis were observed in the granulate matrix adorned withentangled fibers. Neuropils were swollen | 2 |
| Severely injured | Red neurons, pyknotic cells and apoptotic cells were commonly observed.Eosinophilic ghost cells were interspersed in a structureless matrix withdisorganized neuropils and tangling fibers | 3 |
| Deadly injured | Deadly injured. Eosinophilic ghost cells and conspicuous coagulative necrosisprevailed. Several pyknotic cells and red neurons interspersed in the clutteredmatrix, which was filled with vacuoles left by dead neurons | 4 |
